# Global predictions for the risk of establishment of Pierce’s disease of grapevines

**DOI:** 10.1101/2022.05.20.492796

**Authors:** Àlex Giménez-Romero, Javier Galván, Marina Montesinos, Joan Bauzà, Martin Godefroid, Alberto Fereres, José J. Ramasco, Manuel A. Matías, Eduardo Moralejo

## Abstract

The vector-borne bacterium *Xylella fastidiosa* is responsible for Pierce’s disease (PD), a lethal grapevine illness that originated in the Americas. The international plant trade is expanding the geographic range of this pathogen, posing a new threat to viticulture worldwide. To assess the potential incidence of PD, we have built a dynamic epidemiological model based on the response of 36-grapevine varieties to the pathogen in inoculation assays and on the vectors’ distribution when this information is available. Key temperature-driven epidemiological processes, such as PD symptom development and recovery, are mechanistically modelled. Integrating into the model highresolution spatiotemporal climatic data from 1981 onward and different infectivity (*R*_0_) scenarios, we show how the main wine-producing areas thrive mostly in non-risk, transient, or epidemic-risk zones with potentially low growth rates in PD incidence. Epidemic-risk zones with moderate to high growth rates are currently marginal outside the United States. However, a global expansion of epidemic-risk zones coupled with small increments in the disease growth rate is projected for 2050. Our study globally downscales the risk of PD establishment while highlighting the importance of considering climate variability, vector distribution and an invasive criterion in obtaining accurate risk maps to guide policy decision-making in plant health.

---

Emerging plant pathogens and pests are costly both economically and environmentally for society [1–4]. Among valuable crops recurrently affected by emerging diseases, grapevine occupies a remarkable place in the history of plant pathology [5–8]. Nowadays, Pierce’s disease (PD) is considered a potential major threat to winegrowers worldwide [9]. The annual economic burden in California alone has been estimated over $100 million [10], and the disease is a well-recognised limiting factor in the cultivation of *Vitis vinifera* in the southeastern United States [9]. In Europe, despite strict quarantine measures to protect the wine industry (Directive 2000/29/EC), PD has recently been established for the first time in vineyards on the island of Majorca, Spain [11, 12]. This finding alongside the detection of PD in Taiwan [13] has raised concerns about its possible spread to continental Europe and other wine-producing regions worldwide.

The causal agent of PD [14], the bacterium *Xylella fastidiosa* (Xf) [15], is native to the Americas where it also causes vector-borne diseases on many economically important crops, such as citrus, almond, coffee and olive trees [16, 17]. Xf is phylogenetically subdivided into three major monophyletic clades that correspond to the three formally recognised subspecies: *fastidiosa, multiplex* and *pauca*, native from Central, North and South America, respectively [18, 19]. Although as a taxonomic unit Xf infects more than 560 plant species [20], it also shows genetic variation among subspecies and sequence types (STs) in both host specificity and host range [21]. Since 2013, diverse STs of the three subspecies have been detected in Europe mainly associated with crop and ornamental plants [22–24]; among these, the clonal lineage of the subsp. *fastidiosa* responsible for PD (hereafter termed Xf_PD_). The same genetic lineage also causes almond leaf scorch disease in California [25] and Majorca (Spain) [26], where it is widespread in almond plantations and vineyards, affecting more than 23 grape varieties [12].

A key trait in the understanding of Xf’s invasive potential is its capacity of being transmitted non-specifically by xylem sap-feeding insects belonging to sharpshooter leafhoppers (Hemiptera: Cicadellinae) and spittlebugs (Hemiptera: superfamily Cercopidae) [27, 28] – e.g., at least eight species transmit PD in the southeastern United States [29]. Such non-specificity would have facilitated Xf_PD_ invasion after being unwittingly brought to Majorca around 1993 with infected almond cuttings from California and its spread thereafter to grapevines through local populations of the meadow spittlebug, *Philaenus spumarius* [26]. Recently, the role of *P. spumarius* in the transmission of PD in Majorca has been demonstrated [12] and its involvement in epidemic outbreaks in California, previously thought marginal [30, 31], is being revisited [32, 33]. To date, the meadow spittlebug has been confirmed as the major vector in the olive quick decline syndrome, PD and the almond leaf scorch disease outbreaks in Europe [12, 26, 28, 34]; therefore, its geographic distribution should be taken into account when assessing the risk of Xf-related diseases [35].

The tropical origin of Xf subsp. *fastidiosa* already suggests that PD is a thermal-sensitive disease, with the temperature being a range-limiting factor [36, 37]. Thus, the accumulated heat units (i.e., growing-degree days) required to complete the process from Xf_PD_ infection to symptom development is critical to predicting the probability of developing PD acute infections. Conversely, the effect of cold-temperature exposures in the recovery of Xf-infected grapevines is a well-established phenomenon [38–40], limiting the geographic range and damage of chronic PD in vineyards in the United States [9]. Such “winter curing” has been linked to the average *T*_*min*_ of the coldest month, to exposures of extreme-cold temperature for several days, or the accumulation of chilling hours [41]. The dynamics of chronic infections –i.e., those that persist from one year to the next year– is determined by the net balance between the number of new infections during the growing season and of infected plants recovered in winter. Because new infections late in the growing season are more likely to recover during winter than early-season infections, the phenology of the vector has a great influence on the dynamics of chronic infections and PD transmission [30, 42–44].

Several works have attempted to predict the potential geographic range of the subsp. *fastidiosa* [45–47] and other Xf subspecies in Europe [48, 49] and worldwide [47] using bioclimatic correlative species distribution models (SDMs). However, none of these works have explicitly included information on vectors’ distribution or disease dynamics. They hence provide little epidemiological insights into the underlying environmental causes underpinning or limiting a potential invasion. An alternative to overcome these limitations is to develop mechanistic models based on the physiology of the pathogen [50], coupled with epidemiological models that consider the disease dynamics while avoiding the difficulties of including transmission parameters for each of the PD potential vectors.

Risk maps often represent an average snapshot that overlooks interannual climate variability and the effects of climate change as limiting disease factors *per se*. This leads frequently to risk overestimation [51–54]. Increased availability of computational resources to deal with demanding climate databases now makes it possible to fit dynamic epidemiological models that include climate variability at broad spatiotemporal scales. For example, high-resolution satellite based climate data have been employed for testing mechanistic models that relate critical physiological processes of coffee rust with climate variables in past outbreak events [55]. Despite these important advances, no attempt of exploring mechanistic SDM has been performed yet for PD.

In this work, we present a temperature-driven dynamic epidemiological model to infer where PD would have become endemic in different wine regions worldwide from 1981 onward if we forced the introduction of Xf-infected plants. We follow an invasive criterion as defined by Jeger & Bragard [56] to include, as far as we can, key plant, pathogen, and vector parameters and their interactions for estimating the risk of establishment, persistence, and subsequent epidemic development. The model assumes a local Xf_PD_ spatial propagation among plants mediated by the presence of potential vectors. Due to the limited knowledge about the vectors of PD in most wine-growing regions of the world [30], we employ fixed basic reproductive numbers (*R*_0_) in the epidemiological models, except for Europe, where there are precise estimations of climate suitability for the main vector *P. spumarius* [35]. This heuristic approach to obtaining PD risk maps yields results that are consistent with all the relevant data available [45]. It also allows us to quantitatively approximate the current potential growth rate of PD incidence in wine-growing regions under different transmission scenarios, as well as extrapolating the impact of PD by 2050 [57]. By estimating a lower global risk of PD, our study casts doubts on the potential impact predicted for other Xf-related diseases transmitted by *P. spumarius* [49], specially in Europe when vector distribution is taken into account.

## Results

### Thermal requirements to develop PD

We screened a wide spectrum of European grapevine varieties response to Xf_PD_ infection. Overall, 86.1% (*n* = 764) of 886 inoculated plants, comprising 36 varieties and 57 unique scion/rootstock combinations, developed PD symptoms at 16 weeks post-inoculation. European *V. vinifera* varieties exhibited significant differences in their susceptibility to Xf_PD_ (Supplementary Table S1). All varieties, however, showed PD symptoms to some extent, confirming previous field observations of general susceptibility to Xf_PD_ [9, 12, 37]. We also found significant differences in virulence (*χ*^2^ = 68.73, df = 1, *P* = 2.2 ***×*** 10^−16^) between two Xf_PD_ strains isolated from grapevines in Majorca across grapevine varieties (Supplementary Fig. S1). Full details on the results of the inoculation tests are available in Methods, Supplementary Section S1, Supplementary Table S1 and Supplementary Data 1.

The number of symptomatic leaves of each of the inoculated plants was monitored over time. These data were pooled to estimate an average response among grapevine varieties worldwide to thermal accumulation requirements to develop PD under daily fluctuating temperatures. To do this, we introduced the experimental relationship between Xf-growth rate and temperature [38] into the computation of the growing-degree days (Fig. 1A, Methods). The resulting Modified Growing-Degree Days (MGDD) represent a more accurate metric connecting Xf_PD_ growth and symptom development under field conditions (Supplementary Section S2 B). Using survival analysis, we fitted the probabilistic function ℱ (*MGDD*) to develop five or more symptomatic leaves (i.e., assumed as a threshold for chronic infections, see Supplementary Section S1). We thus found a minimum window of *MGDD* = 528 to develop chronic infections (*var*. Tempranillo), about 975 for a Kaplan-Meier median estimate, while a cumulative *MGDD >* 1159 indicated over 90% probability within a growing season (red curve in Fig. 1C and Methods).

**FIG. 1.**
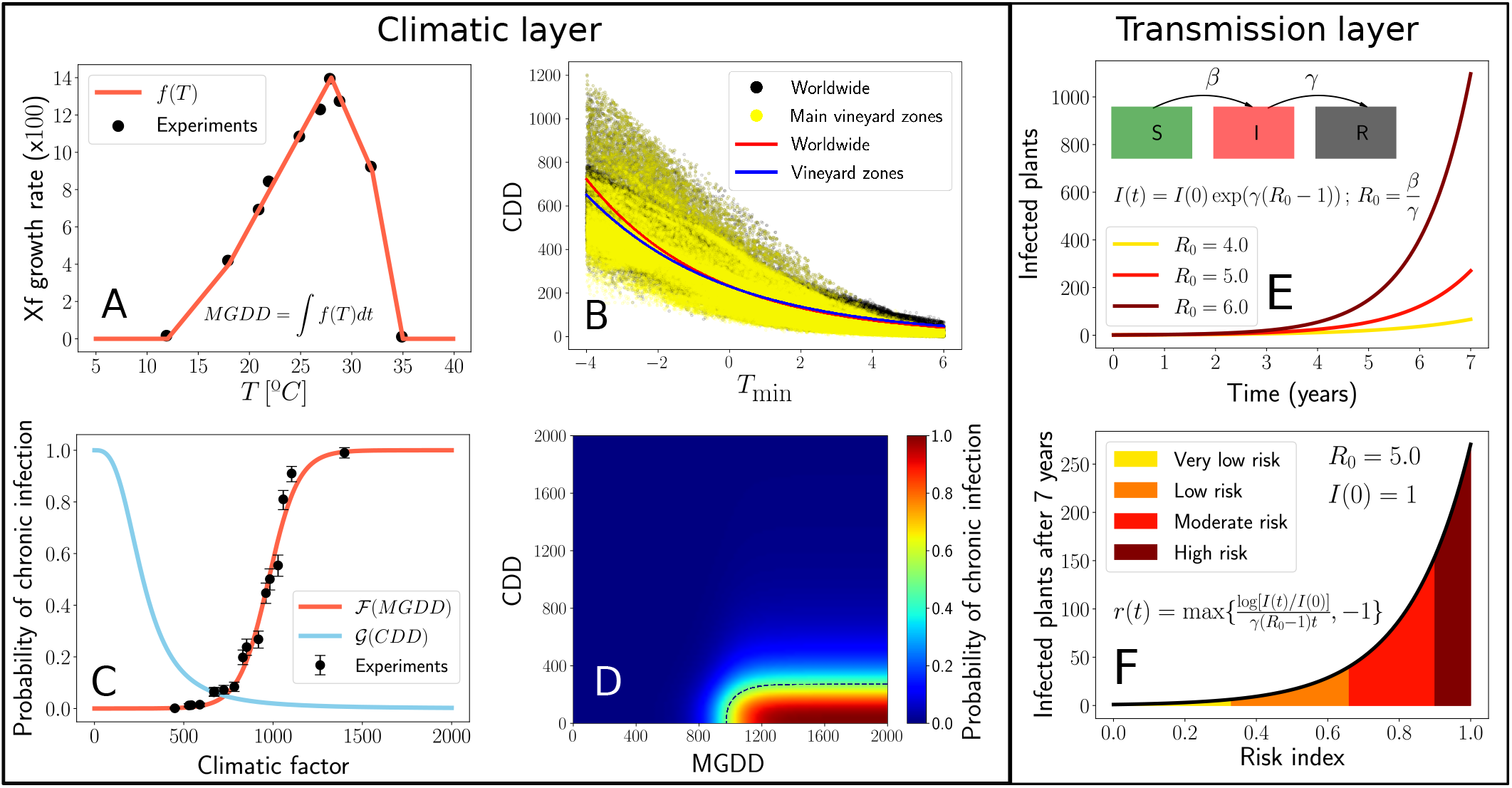
Climatic and transmission layers composing the epidemiological model. (**A**) *MGDD* profile fitted to the data of Xf growth rate in Feil & Purcell 2001 [38]. The original Arrhenius plot, log *k vs*. 1*/T*, in kelvin was converted to a linear dependence in Celsius temperature *T* (Supplementary Section S2 A). (**B**) Correlation between *CDD* and the minimum average temperature of the coldest month between 1981 and 2019. Plotted black dots (worldwide) and yellow dots (main wine producing zones) depict data distributed around all the world at resolution 0.1° ***×*** 0.1° (6,487,200 cells) retrieved from ERA5-Land dataset. Red solid line depicts the fitted exponential function for worldwide data and blue solid line for main vineyards zones. (**C**) Nonlinear relationship between *MGDD* (red line) and *CDD* (blue line) with the likelihood of developing chronic infections. Black dots depict the accumulated proportion of plants across 36 grapevine varieties showing five or more symptomatic leaves at 15 *MGDD* levels (see Supplementary Information). Vertical bars are the 95% CI. (**D**) Combined ranges of *MGDD* and *CDD* on the likelihood of developing chronic infection. (**E**) Transmission layer in the dynamic equation (1) of the SIR compartmental model. (**F**) Relationship between the exponential growth of the number of infected plants with the risk index and their ranks.

We then investigated the probability of disease recovery by exposure to cold temperatures. However, unlike the approach followed to relate MGDDs to Xf_PD_ growth and symptom development, there are sparse experimental data to lean on. Following the work of Lieth et al.[39] and an inverse analogy with the MGDDs, we assume that cold temperatures contribute to the decrease of the bacterial population *in planta*. Thus, the accumulation of cold degree-days (CDD) is expected to be a proper metric to describe the probability of recovery of infected plants. To account for the effect of winter curing, we related the accumulation of cold degree-days (CDD with base temperature = 6 °C, i.e., 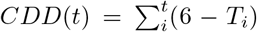 for *T*_*i*_ ≤ 6°C) with the average minimum temperature of the coldest month, *T*_*min*_, isolines in the United States proposed by Purcell (available in [41]) as reference zones where PD is rare (*T*_*min*_ between − 1.1 °C and 1.7 °C), occasional (1.7 − 4.5°C) and severe (*>* 4.5°C) (Fig. 1B, Methods). This allows to relate accumulated CDD to the probability of chronic infection (i.e. the complementary probability of disease recovery) using a generalised sigmoidal function, 𝒢 (*CDD*) (see blue curve in Fig. 1C and Methods).

### MGDD/CDD distribution maps

MGDDs were used to compute annual risk maps of developing PD during summer for the period 1981-2019. The resulting averaged map identifies all known areas with a recent history of severe PD in the United States corresponding to ℱ (*MGDD*) *>* 90% (i.e. high-risk), such as the Gulf Coast states (Texas, Alabama, Mississippi, Louisiana, Florida), Georgia and Southern California sites (e.g., Temecula Valley) (Fig. 2*A*), while captures areas with a steep gradation of disease endemicity in the north coast of California ℱ (*MGDD >* 50%). Overall, more than 95% of confirmed PD sites (*n* = 155) in the United States (Supplementary Data S2) fall in grid cells with ℱ (*MGDD*) *>* 50%.

**FIG. 2.**
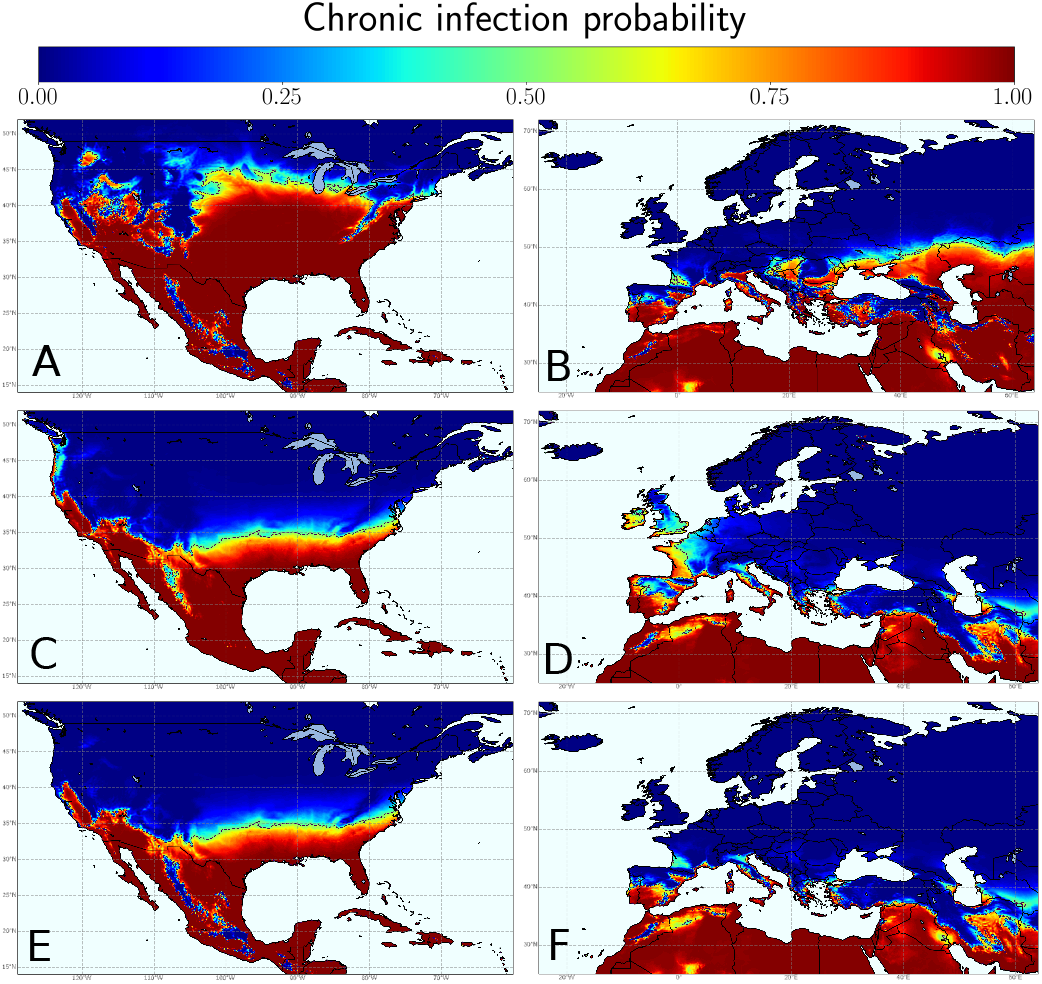
Average thermal-dependent maps for Pierce’s disease (PD) development and recovery in North America and Europe. PD development during the growing season based on average ℱ (*MGDD*) estimations between 1981 and 2019 in North America (**A**) and Europe (**B**) derived from the results of the inoculation experiments on 36 grapevine varieties. Large differences in the areal extension with favourable MGDDs can be observed between the United States and Europe. The winter curing effect is reflected in the distribution of the average 𝒢 (*CDD*) for the 1981-2019 period in the United States (**C**) and Europe (**D**). A snapshot of the temperature-driven probability of chronic infection averaged in the 1981-2019 period is obtained from the join effect of MGDD and CDD in North America (**E**) and Europe (**F**). Warmer colours indicate more favourable conditions for chronic PD and the dashed line highlights the threshold of chronic infection probability being 0.5.

The average MGDD-projected map for Europe during 1981-2019 spots a high-risk for the coast, islands and major river valleys of the Mediterranean Basin, southern Spain, the Atlantic coast from Gibraltar to Oporto, and continental areas of central and southeast Europe (Fig. 2B). Of these, however, only some Mediterranean islands such as Cyprus and Crete show ℱ (*MGDD*) *>* 99% comparable to areas with high disease incidence in the Gulf Coast states of the United States and California. Almost all the Atlantic coast from Oporto (Portugal) to Denmark are below suitable *MGDD*, with an important exception in the Garonne river basin in France (Bordeaux Area) with low to moderate *MGDD* (Fig. 2B).

Fig. 2A shows how the area with high-risk *MGDD* values extends further north of the current known PD distribution in the southeastern United States, suggesting that winter temperatures limit the expansion of PD northwards [9]. A comparison between MGDD and CDD maps (Fig. 2A vs. Fig. 2C, Fig. 2E) further supports the idea that winter curing is restricting PD northward migration from the southeastern United States. However, consistent with growing concern among Midwest states winegrowers on PD northward migration led by climate change [58], we found a mean increase of 0.12% y^−1^ in the areal extent with *CDD <* 306 (∼ *T*_min_ *<* −1.1 °C) since 1981 comprising land areas between 103^*o*^W and 70^*o*^W of the United States (Supplementary Fig. S4). Such an upward trend corresponds to 5090 km^2^ y^−1^ in the potential northward expansion of PD due to climate change, and an accumulation of ∼ 193,420 km^2^ of new areas at risk since 1981.

High-*CDD* values would also be expected to restrict the potential PD colonisation in continental Europe (Fig. 2D). Unlike North America, the East-West distribution of major European mountain ranges together with the warming effect of the Gulf Stream decrease the likelihood of cold winter spells reaching the western Mediterranean coast. 𝒢 (*CDD*) between 100% and 95% (i.e., recovery probability *<* 5% – low winter curing) are mostly prevalent below 40^*o*^N latitude in southwest Iberian Peninsula and Mediterranean islands and coastlands (*<* 50 km away). Above 40^*o*^N latitudes, *CDD <* 100 are encountered mainly in the Atlantic coast and Mediterranean coast and islands (Fig. 2D). In contrast, continental central and southeast Europe show high *CDD* values likely preventing Xf_PD_ winter survival on infected grapevines.

In Fig. 2E-F, we show the average climatic suitability for PD establishment only from the mechanistic relation between Xf_PD_ and temperature. Although all areas with current Xf_PD_-related outbreaks are identified, risk predictions based only on the combination of MGDD and CDD could lead to overestimations, as this approach overlooks disease transmission dynamics and climate interannual variability. These flaws are corrected in the complete model below.

### Disease Model

Our mathematical model is based on a compartmental SIR framework in which plants can be in susceptible, infectious/infected, or eliminated states with respect to disease. Moreover, the model accounts for the thermal requirements to develop PD by incorporating both ℱ (*MGDD*) and 𝒢 (*CDD*) functions. The global computation of MGDD and CDD is based on hourly temperature data from ERA5-Land dataset [59] at 0.1^*o*^ spatial resolution (Methods), which defines the spatial units of our model (i.e. cells of 0.1^*o*^ ***×*** 0.1^*o*^). Within the cells, we approximate vector-plant disease transmission with an effective plant-to-plant transmission rate *β* (Supplementary Section S4), as done in previous Xf-related diseases [60]. Since our aim is to estimate the risk of establishment, we assume the presence of an initial population of infected plants in every cell and study its potential growth or decrease, while disease propagation among cells is not considered. The SIR models show an exponential growth/decrease regime for the infected population in the very early stages. This behaviour is governed by the basic reproductive number (*R*_0_ = *β*/*γ*), which in a mean-field approximation defines the onset of an outbreak (Fig. 1E, Supplementary Section S2C), where γ is the mortality rate of PD infected vines fixed at 1/5 years^−1^ [25]. Different values of *β*, and consequently of *R*_0_, will be explored throughout the study.

To address if new infections will become chronic, we apply the combination of ℱ (*MGDD*) and 𝒢 (*CDD*) functions (Fig. 1D) to the output of the transmission layer at the end of each year. Throughout time (years), *R*_0_ must be greater than 1 for the size of outbreak to trend upwards. The model assumes a homogeneous distribution of the vector among the cells (same *β* and then same *R*_0_) except for Europe, where information on the spatial distribution of *P. spumarius* is available (see Methods). Altogether, the equation representing the evolution of the number of chronic infections in each cell *j* at time *t* is written as

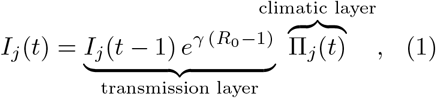

where *I*(*t* − 1) is the number of chronic infections in the previous year and П_*j*_(*t*) = ℱ (*MGDD*) · 𝒢 (*CDD*) is the climatic layer, which modulates the growth term and describes the cumulative probability of new infections becoming chronic. To account for a non-homogeneous vector distribution, a spatial dependent *R*_0_(*j*) is incorporated into the model by considering the product of the homogeneous *R*_0_ and the spatially-dependent climate suitability for vectors (Supplementary Section S2 F).

### Modelling risk

To compute the epidemic-risk maps, we carried out a simple simulation summarised in three steps: (i) at the initial condition setting, each cell is seeded with a single infected plant, *I*(0) = 1; (ii) the simulation runs for a given period, *τ*, and the incidence at each time-step is estimated following Eq. (1); finally, (iii) the risk index *r*_*j*_(*τ*) is calculated from the final number of infected plants at cell *j* as

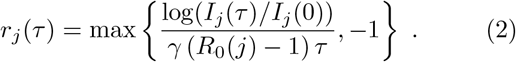

*r*_*j*_(*τ*) implicitly delimits three differential risk zones in the maps: 1) an epidemic-risk zone (*r*_*j*_(*τ*) *>* 0), where PD can become theoretically established; 2) a transition zone, which can be arbitrarily defined by the area delimited by the lines *I*(*τ*) ≥ 10 *I*(0) and *I*(*τ*) ≤ 0.05 *I*(0). In these zones, the probability of PD becoming established ranges between 0 and 1 and the exponential growth rate oscillates around *r*_*j*_(*τ*) = 0; and finally, 3) a non-risk zone characterised by an exponential decrease in the number of infected vines. Our risk index measures, therefore, the relative initial exponential growth rate of the population of infected plants respect to the maximum growth, *r*(*τ*) = 1, which serves to rank the epidemic-risk zones in high (*>* 0.9), moderate (0.66 − 0.9), low (0.33 − 0.66) and very low (∼ 0.1 − 0.33) risks (Fig. 1F, Supplementary Section S2 E).

### Model calibration and validation

To attempt a rough estimate of the *R*_0_ parameter in the United States, assuming a uniform spatial distribution of the vectors, we ran several model simulations validating the spatiotemporal distribution of PD from data collected from publications between 2001 and 2015 [19, 39, 41, 61– 65]. We found *R*_0_ = 8 as the optimal parameter for maximising the area under a ROC curve (Supplementary Fig. S5). In general, our model returns an accuracy of more than 80%, except for 2006, due to data retrieved from a study in the Piedmont region in the USA at the limit of the transient-risk zone (Supplementary Fig. S7 and Table II). A different approach was followed to estimate *R*_0_ for Europe given that PD is only present in Majorca and hence spatiotemporal data on the PD distribution is not available. First, we estimated the transmission rate of the main European vector *P. spumarius* from the well-studied disease progress curve of the almond leaf scorch epidemic in Majorca. Then, using the known mortality rate of PD-infected vines *γ* ∼ 1*/*5 years^−1^ and the inferred transmission rate, *β* = 0.8 years^−1^, the basic reproduction number for PD in Majorca yields *R*_0_ = *β/γ* = 4. Finally, using data on the climate suitability of the vector in Majorca, *v* = 0.8, and inverting the relation *R*_0_(*j*) = *R*_0_ *v*(*j*), we estimated *R*_0_ = 4*/*0.8 = 5 as a baseline scenario for PD transmission in Europe (Supplementary Section S2 D). This figure is not intended to be an exact estimate of *R*_0_ but rather an average reference in our model in agreement with the lesser abundance of vectors relative to the United States. Furthermore, since there is no information on the distribution of the potential vectors and no PD distribution data to calibrate, we also used a conservative *R*_0_ = 5 scenario for the rest of the world.

**TABLE I.**
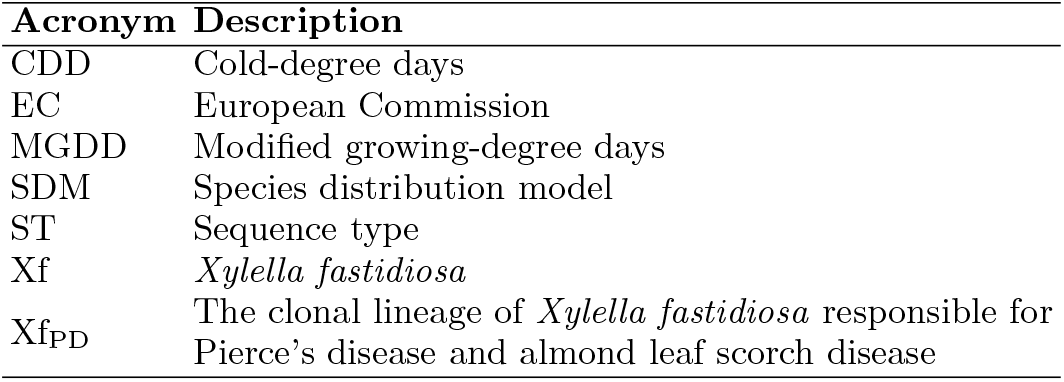
Abbreviations used in the text

**TABLE II.**
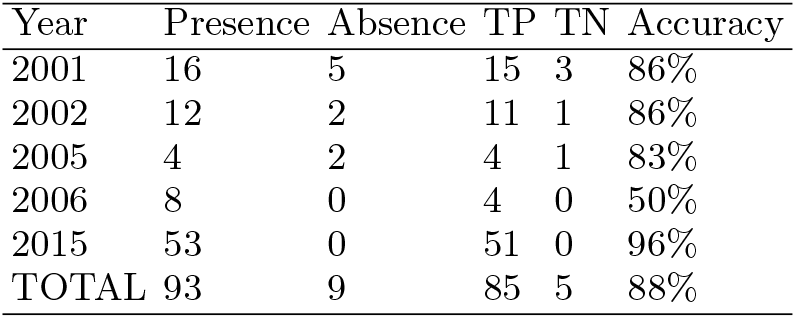
**Validation of model predictions**. The items are locations where PD was present or absent. TP corresponds to true positives and TN to true negatives according to our model with *R*_0_ = 8.

### PD global risk

Currently, PD is mainly restricted to the American continent with some unrelated introductions of Xf_PD_ to Taiwan and Majorca (Spain) from the United States [12, 13]. To probe the climatic suitability of other potential regions, we projected the dynamic model to main winegrowing regions in the Northern Hemisphere (United States, Europe and China) and Southern Hemi-sphere (Chile, Argentina, South Africa, Australia and New Zealand)(Fig. 3A-E).

**FIG. 3.**
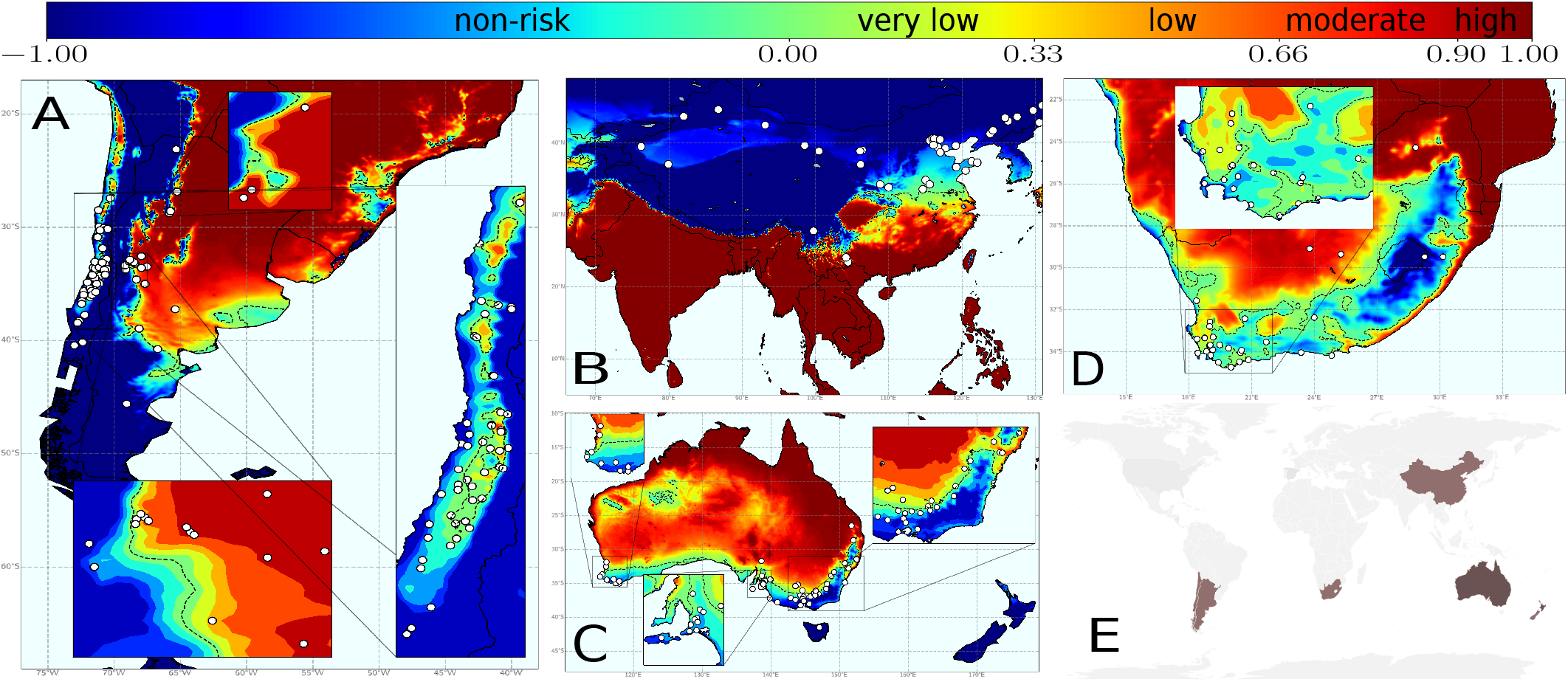
Climate-driven risk maps for PD establishment in main viticulture regions worldwide under a baseline *R*_0_ = 5 scenario. White dots indicate the main vineyard areas in the wine-growing regions of China and the Southern Hemisphere. (**A**) Chile and Argentina; (**B**) Asia with special attention to China; (**C**) Australia and New Zealand (wine areas are not marked as the whole country is without risk); and (**D**) South Africa. (**E**) Global distribution of main wine-producing areas analysed. The risk index *r*_*j*_(*τ*), express the relative exponential growth rate of the disease incidence, and was scaled from 0.1 to 1 and ranked as very low (0.10-0.33), low (0.33-0.66), moderate (0.66-0.90) and high (*>* 0.90).

Emerging wine-producing areas in China are predominantly located in non-risk zones, whereas only some vineyards in the Henan province fall in transition risk zones (Fig. 3B). In Europe, 92.1% of the territory is at nonrisk and 6.1% is included in the epidemic-risk zone, with only 1.9% showing a high-risk index and 1.5% a moderate risk (Supplementary Table S2). The model also reveals a progressive transition from areas without risk (*r*(*t*) *<* 0) before 1990 to epidemic-risk zones potentially with low-risk indexes by 2019 ([57], see Movies), mainly affecting the basins of the rivers Po in Italy, Garonne and Rhone (France) and Douro/Duero (Portugal and Spain). This represents a mean increase of 0.21% y^−1^ in the risk-epidemic zone, a rate 3.5-times greater than that of the eastern United States, which could increase the likelihood of PD establishment in Europe in the coming decades. In the United States most states around the Gulf Coast show high risk-indexes, whereas on the west coast, around 37.5% of California surface is suitable for epidemics with high growth rates (Supplementary Table S3).

In the Southern Hemisphere, vineyards at non-risk and transient epidemic-risk zones predominate – e.g., non-risk in New Zealand and Tasmania (Fig. 3C). Risk indexes in areas where PD can become established (*r*(*t*) *>* 0) range from very low to low for most coastal vineyards in Australia (west, south and east) with some more suitable conditions in the interior of New South Wales, Greater Perth and Queensland (Fig. 3C); a general very-low or low-risk indexes are predicted in the Western Cape in South Africa (Fig. 3D); overall very-low but localised low to moderate risk indexes in some areas in Chile; and low to moderate growth of the number of infected vines in most of Argentina; being this the wine-growing country with the highest risk (Fig. 3A). Detailed information on areas with non-risk, transient risk and risk indexes (i.e., disease-incidence growth rates) in areas with the potential risk of establishment by country and regions is provided in Supplementary Table S4 and Supplementary Data S3.

Within epidemic-risk zones, risk indexes may vary if any of the epidemiological parameters governing transmission change. As expected, *I*(*t*) *< I*(0) boundaries increasingly displace to northern latitudes in the US and Europe under higher transmission scenarios, increasing the risk-epidemic zones significantly (Fig. 4A-F). The line representing the outbreak extinction i.e, the non-risk zone *r*(*t*) *<* −0.09, in the validated *R*_0_ = 8 scenario for the United States, falls at some distance above the isoline *T*_min_ =− 1.1^*o*^C in comparison to the *R*_0_ = 5 scenario (Fig. 4C vs Fig. 4A and [57], Movies). This distribution pattern holds and moves slightly northward over time in parallel to global warming, although the trend of PD latitudinal change is moderated by high-*CDD* values (i.e. cold accumulation). In addition, the disease extension also declines due to *CDD* interannual fluctuations in the simulations. Cold waves periodically occur that reach latitudes close to the Gulf, such as those that occurred in 1983, 1993, 1995, 2000, 2009 and 2013 (Movies at [57]), thus preventing PD expansion northward. The magnitude of this decrease is revealed after comparing the average annual increase of the areas between *r*(*t*) *>* 0 and *CDD <* 314 lines. From 1981 to 2019, the area with risk *r*(*t*) *>* 0 increased at a rate of 0.05% (y^−1^), while that of *CDD <* 314 by 0.12% y^−1^, an important difference not explained alone by *CDD*s without considering climate fluctuations (Supplementary Fig. S4).

**FIG. 4.**
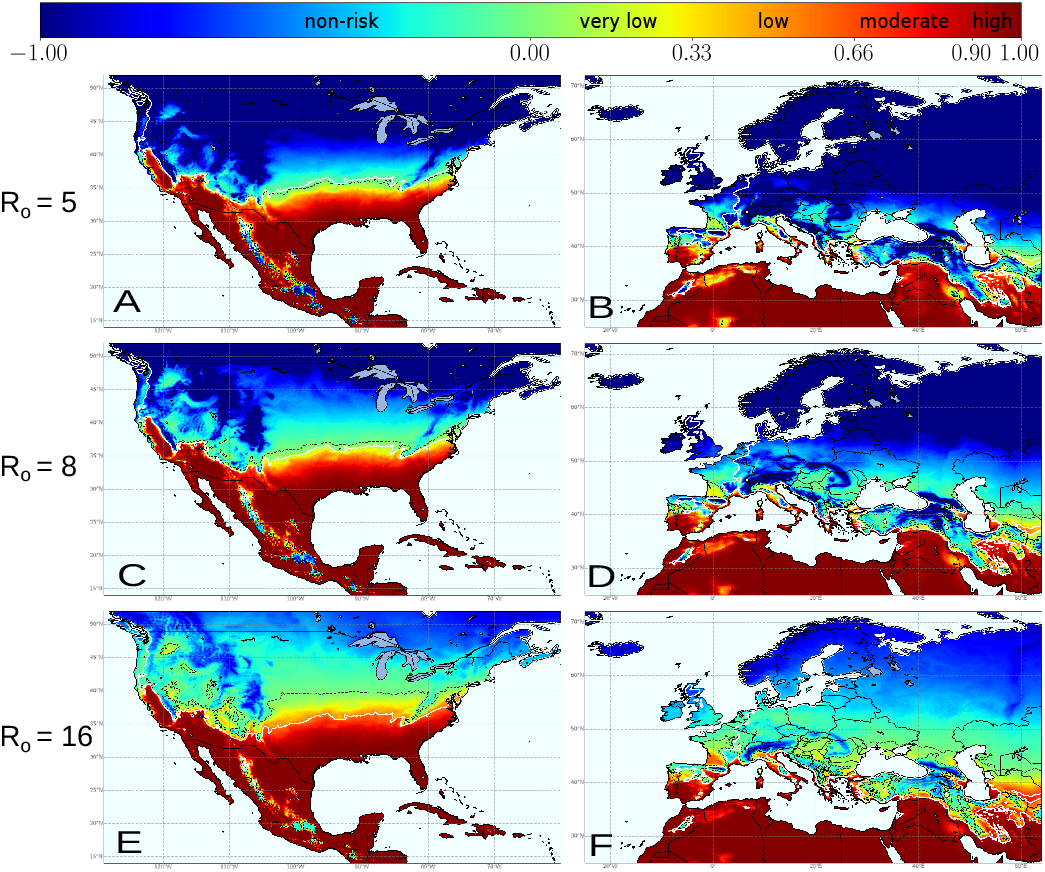
Climate-driven dynamic-model simulations for PD establishment from 1981 to 2019 under different *R*_0_ scenarios with a spatially homogeneous vector distribution. For comparison, the baseline scenario with a *R*_0_ = 5 for Europe is projected to North America **(A)** capturing to some extent the distribution and severity of PD in that continent. In Europe (**B**) high-risk areas (i.e., *r*_*j*_ (*τ*) *>* 0.90) are restricted to coastal Mediterranean and the south of the Iberian Peninsula; black dash line separate areas with *r*(0) *>* 0 where theoretically PD can thrive. Under higher *R*_0_ scenarios, *R*_0_ = 8 for North America (**C**) and Europe (**D**), the dash lines tend to separate from isoline *T*_min_ = − 1.1°C (white line); and even more in extreme transmission pressure *R*_0_ = 16 for North America (**E**) and Europe (**F**).

### PD risk projections for 2050

Global shifts in the risk index *r*_*j*_(*τ*) between 2019 and those projected for 2050 were calculated under the same baseline scenario (Fig. 5A-F, Methods). Our simulation shows a generalised increasing trend mainly due to shifts from transition zones to epidemic-risk zones with very low or low-risk indexes in the main wine-growing regions, except for the United States. Here the epidemic-risk zone would increase by 12.8% with the greater increments in the high-risk index category (22.7%) and a decrease in the transition zones (Supplementary Table S5). Much less surface would fall in the epidemic-risk zone in Europe (8.6%) compared to the United States (36.5%); however, the epidemic-risk zone would expand by 40.0% with respect to 2020, a rate over three times greater than in the United States (Supplementary Table S6). Such increases are due to the emergence of previously unaffected areas in 2020 evolving to epidemic-risk zones by 2050, and epidemic growth-rate increases in already epidemic-risk zones in 12 of 42 countries (Supplementary Table S2). Among these 12 countries, however, there is substantial variation in the risk index increments within epidemic-risk zones with respect to 2019 (Supplementary Table S6). While non-risk zones still cover 87.6% of Europe land area, epidemic-risk zones with high-risk indexes are expected to be almost two-fold higher than that of 2019, comprising 3.2% of Europe (Table III).

**FIG. 5.**
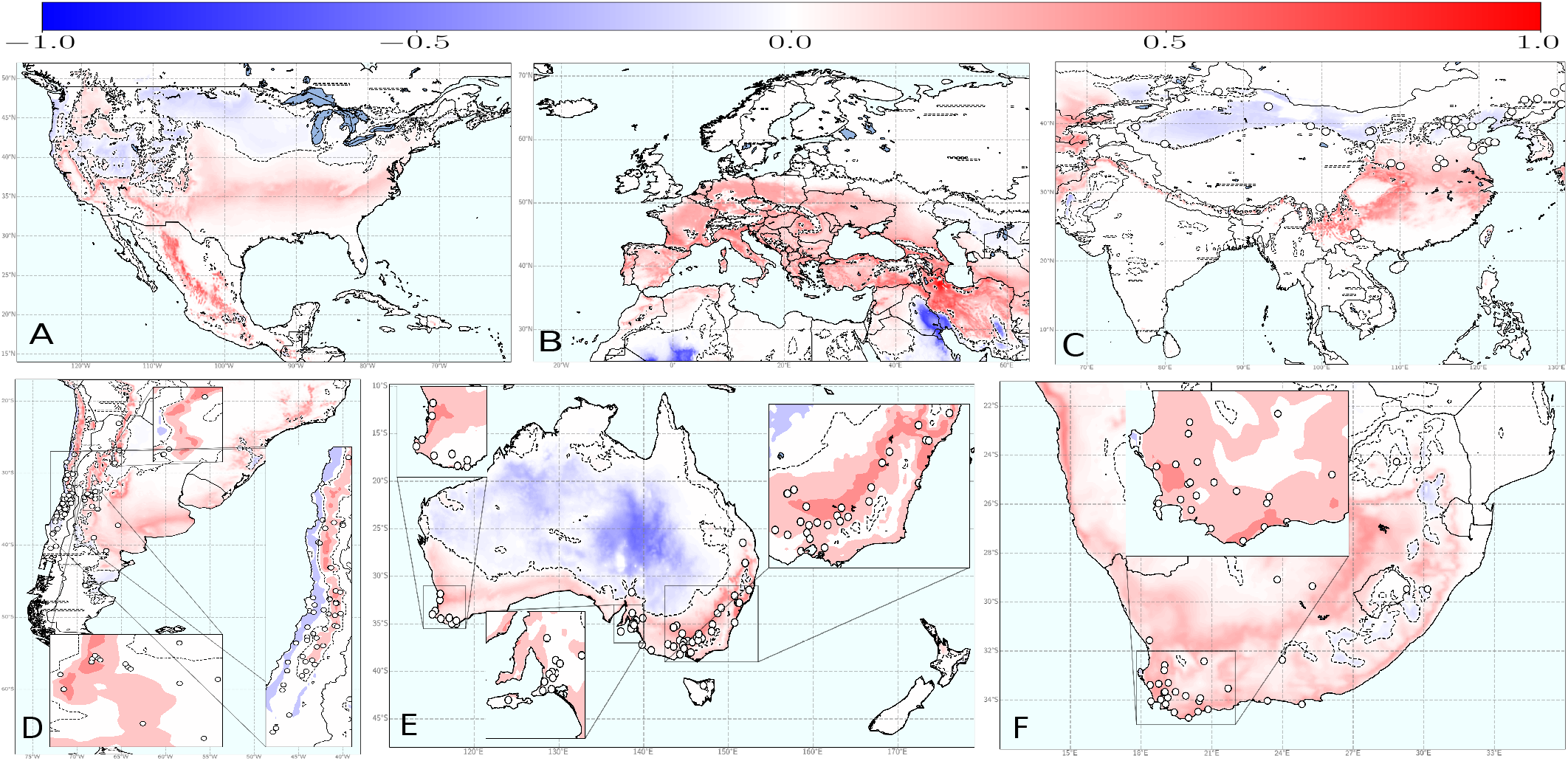
Global shifts in PD risk index (*r*_*j*_ (*τ*)) from 2020 to 2050. To build the maps, we have assumed a spatial homogeneous vector distribution and a *R*_0_ = 5 scenario, except for the United States where a *R*_0_ = 8 has been used in the model simulations.**(A)** North America;**(B)** Europe;**(C)** Asia; **(D)** South America; **(E)** Australia and New Zealand; and **(F)** South Africa. Risk-index increases are in red and decreases in blue. Dash line represents the spatial threshold where *r*_*j*_ (*τ*) difference changes from negative to positive.

**TABLE III.**
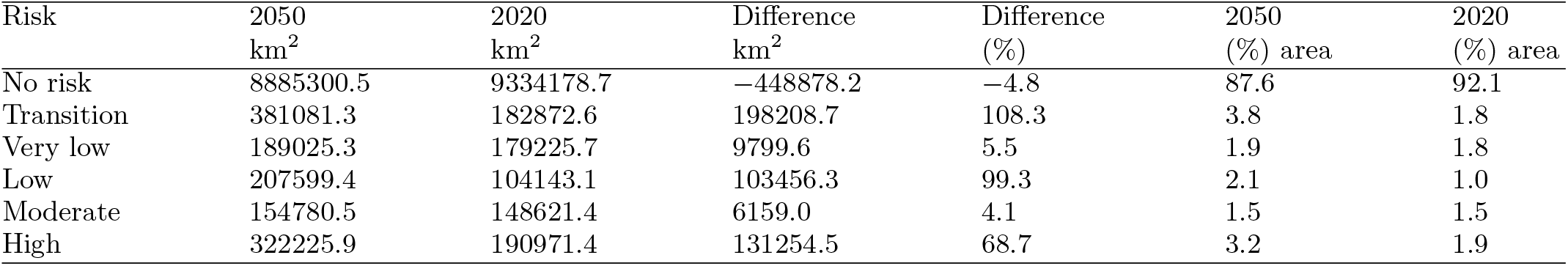
**Shifts in risk areas for Pierce’s disease in Europe projected for 2050 under a** *R*_0_ **= 5 scenario**. The model was run assuming the same homogeneous spatial distribution of the vector for the whole period.

| Risk | 2050<br>km<sup>2</sup> | 2020<br>km<sup>2</sup> | Difference<br>km<sup>2</sup> | Difference<br>(%) | 2050<br>(%) area | 2020<br>(%) area |
| --- | --- | --- | --- | --- | --- | --- |
| No risk | 8885300.5 | 9334178.7 | -448878.2 | -4.8 | 87.6 | 92.1 |
| Transition | 381081.3 | 182872.6 | 198208.7 | 108.3 | 3.8 | 1.8 |
| Very low | 189025.3 | 179225.7 | 9799.6 | 5.5 | 1.9 | 1.8 |
| Low | 207599.4 | 104143.1 | 103456.3 | 99.3 | 2.1 | 1.0 |
| Moderate | 154780.5 | 148621.4 | 6159.0 | 4.1 | 1.5 | 1.5 |
| High | 322225.9 | 190971.4 | 131254.5 | 68.7 | 3.2 | 1.9 |

### Risk based on vector information

So far, we have ignored the distribution of known and potential vector species due to their large number in the Americas and the limited quantitative information generally available. In the case of Europe, given the prevalence of *P. spumarius* as a potential vector and its wide distribution, we added a vector layer in a spatially dependent 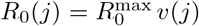, where *v*(*j*) is the climatic suitability for the vector (Methods), *v* = 1 implies optimal climatic conditions for the vector with no constraints for the population size, while *v* = 0 implies unsuitable climatic conditions and its absence (Supplementary Fig. S8). According to the model, no part of Europe shows a high-risk index and barely 0.34% of the territory falls in areas with potential moderate exponential growth rates in disease incidence (Supplementary Table S7). Irrespective of vineyard distribution, we estimated that PD could potentially become established (i.e., *r*(*t*) *>* 0) at a maximum of 3.1% of the territory, while the area at moderate-risk index would be around 5-times lesser than the model without the vector’s climate suitability layer, this latter more in consonance with other proposed risk maps [45, 46]. Such differences in the projected risks are mainly concentrated in the warmest and driest Mediterranean regions and are due to uncertainties concerning temperature-humidity interactions in the ecology of the vector [35].

### Combining vineyard land cover across Europe with the model output

When we integrate to the analysis a layer of vineyards surface from Corine-Land-Cover, we find that PD could potentially become established (i.e., *r*(*t*) *>* 0.075) in 22.3% of the vineyards in Europe. However, no vineyard is at epidemic-risk zones with a high-risk index and only 2.9% of vineyard surface is at moderate risk (Supplementary Table S8). The areas with the highest risk index (*r*(*t*) between 0.70 and 0.88) are mainly located in the Mediterranean islands of Crete, Cyprus and the Balearic Islands or at pronounced peninsulas like Apulia (Italy) and Peloponnese (Greece) in the continent (Fig. 6A and Supplementary Table S8). Most vineyards are at non-risk (42.1%) and another 35.6% are located in transition zones with presently non-risk but where Xf_PD_ could become established in the next decades causing some sporadic outbreaks. All known areas where Xf is well-established in Europe (e.g. Apulia, Corsica, Balearic Islands, Region of Provence-Alpes Côte d’Azur (French Riviera), Alicante) are in the 96th percentile of the tracked sites, validating the strength of our mechanistic, non-correlative PD model predictions (test in [57]). In Supplementary Data S4 and Supplementary Table S8, we provide full details of the total vineyard areas currently at risk for each country and region.

**FIG. 6.**
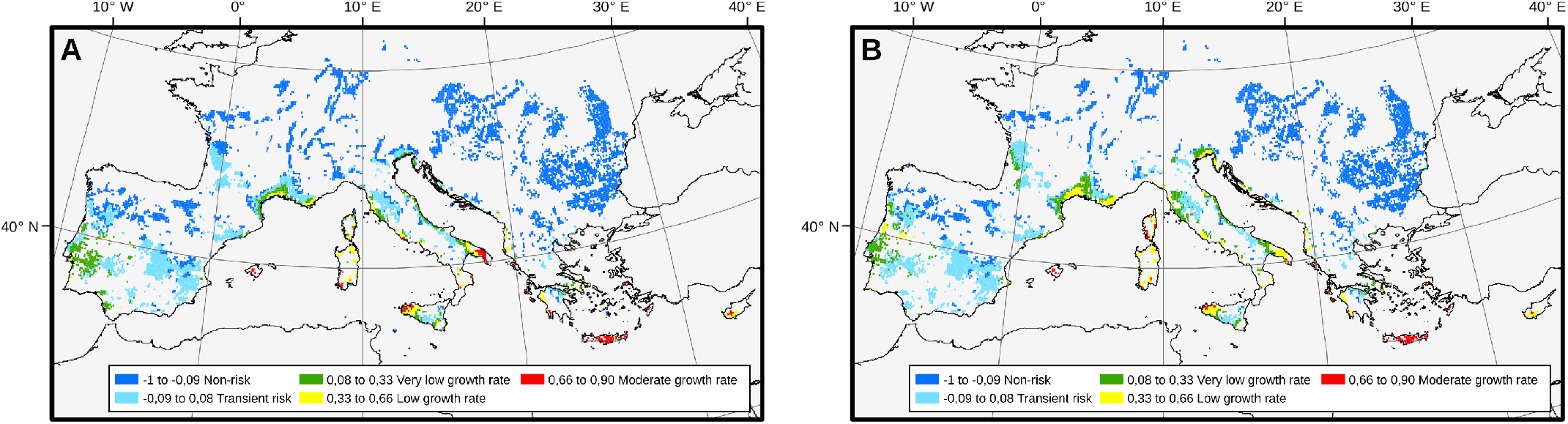
Intersection between Corine-land-cover vineyard distribution map and PD-risk maps for 2020 and 2050. Data were obtained from Corine-land-cover (2018) and the layer of climatic suitability for*P. spumarius* in Europe from [35]. The surface of the vineyard contour has been enlarged to improve the visualisation of the risk zones and disease-incidence growth-rate ranks. **(A)** PD risk map for 2019 and its projection for 2050 **(B)**. Blue colours represent non-risk zones and transient risk zones for chronic PD (*R*_0_ *<* 1). The 2050 map shows some contraction of epidemic-risk zones with moderate risk indexes in Mediterranean islands and Apulia as the climate becomes hotter and dryer.

Our model with climate and vector distribution projections for 2050 indicates a 55.8% increase of the epidemic-risk zone in Europe (Fig. 6B). This increment would be mainly due to the extension of epidemic-risk zones with very low and low-risk indexes. However, within the epidemic-risk zones, the areas with moderate risk indexes would decrease from (114,925 ha in 2020 to 43,114 ha in 2050, and no vineyards would be at high risk (Fig. 6B; see Supplementary Table S9 and Supplementary Data S4). Counterintuitively, our model indicates a substantial increase of the area where PD could establish and become endemic for 2050, but a moderate decline in those areas where crop damage could be expected to be significant (e.g. Balearic Islands, Crete, Cyprus, Apulia).

## Discussion

We introduce an epidemic approach to assessing the risk of establishment of PD in vines. The model includes the dynamics of the infected-host population, which enables estimating the initial exponential growth/decrease rate of the disease incidence. Unlike SDM correlative studies, Bayesian or, in general, machine learning black-box approaches, our model goes beyond by providing a mechanistic framework and thus explanatory power. In addition, it is flexible enough to simulate different climate and transmission scenarios, allowing, for instance, the incorporation of the information on the spatial distribution of the vector. We present comprehensive maps of the global PD risk derived from simulations of our mechanistic model using high-resolution climate data. A web page is included, showing results of simulations with different parameters to estimate the risk of PD anywhere [57].

Temperature rules key physiological processes in ectothermic organisms involved in PD and thus determines the ranges of thermal limitation in which they can thrive [52]. Xf_PD_ multiplication and survival within vine xylem vessels not only characterise PD, but also determine the bacterial population dynamics [38]. PD symptom development can be therefore characterised as a thermal-dependent continuous process within the range of cardinal Xf_PD_-growth temperatures [53]. Metrics of thermal integrals (MGDD) based on robust experimental data provide a reliable predictor of climatic suitability and the probability of developing PD during the summer, whereas CDD account for the effect of cold-temperature exposure in the recovery of the infected plant. This dual effect of MGDD and CDD on the demography of infected plants shapes in our model the impact of climate variability on the epidemic dynamics in the early stages of invasion (Fig. 1D). Because the physiological basis of the plant-Xf interaction leading to symptoms development is poorly understood, we caution that other environmental factors, such as drought, nutrient status or crop management may modulate symptom expression and hence add an error in the MGDD parameter not measured in this work. Nonetheless, we deem the error range would be smaller than the differences in *MGDD* among varieties (i.e., regional differences) and smaller than the interannual *MGDD* oscillations in most locations.

Knowledge of insect distribution is crucial for predicting epidemic outbreaks of endemic diseases, as well as the risk of invasion by emerging vector-borne pathogens ([56, 66], (cf. [49])). Given the great diversity of known and potential vector species that can transmit PD [30], it has not been possible to include in the model each one of the particular vectors of each region. Therefore, when evaluating the risk of PD on a global scale, we have considered a homogeneous spatial distribution of the vector (fixed *R*_0_), except in Europe where there is information on the main vector (Supplementary Fig. S8). As expected, the European case shows how models that assume a homogeneous spatial distribution of the vector generally produce epidemic risk zones with higher risk indices than models that include a heterogeneous spatial distribution (Supplementary Table S2 vs. Supplementary Table S7). This lack of vector information is one of the main reasons why vector-borne plant diseases are generally overestimated in risk assessments.

Involuntary risk overestimation may stem from other additional sources too. The use of mean data as inputs in epidemiological models can lead to biased results when response functions are nonlinear and climate variability is not accounted for [53]. In this study, we present experimental evidence of a non-linear relationship between MGDD and PD symptom development and indirect empirical evidence of a non-linear relationship between CDD and PD recovery (Supplementary Fig. S9). Such a nonlinear response consequently has a great impact on reducing the risk of PD establishment and steeping the spatial gradients in risk maps (Figs. 4 and 6). Moreover, MGDD and CDD might help to explain why disease pressure is much higher in the southeastern United States than in California and Europe (Figs. 2 and 4) or, for example, earlier reports of PD outbreaks in Kosovo [67]. Cooler summer-nights in California and a shorter growing season compared to those found in the Gulf states in the southeastern United States explain the difference in the accumulated *MGDD* for both areas. In the case of Kosovo, *CDD* values above certain thresholds could have led to the extinction of incipient outbreaks driven by several years with *MGDD* in the conducive range of PD (Fig. 2).

Our PD risk map for Europe confirms earlier correlative SDM-derived predictions for the subsp. *fastidiosa* [45]. Both approaches make congruent predictions on PD potential distribution, providing convergent lines of independent evidence of climate suitability. However, our risk maps go further by incorporating in the epidemic-risk zones information on the relative exponential growth rates in the potential disease incidence. By considering different scenarios, this information could be used to assess intervention policies. In general terms, the epidemic-risk map including vector’s information indicates a low risk for chronic PD. Only ∼ 0.34% of European vineyard surface, mainly located in Cyprus, Crete, Sardinia, part of Sicily and the Balearic Islands, meet climatic conditions for PD to become endemic and cause significant damage (Supplementary Table S7 and Supplementary Data S4). Other regions such as Bordeaux, Portugal, Rhône Valley, the Veneto region, would be included in epidemic-risk zones but with very low to low exponential growth rates in disease incidence. By contrast, notorious winegrowing regions in Spain (e.g., Rioja, Ribera del Duero), France (e.g., Burgundy) and Italy (e.g., Piedmont) currently fall within areas considered as non-risk zones, transient-epidemic zones or epidemic-risk zones with very-low risk indexes (Fig. 6).

The dynamic nature of the simulation outputs already points to a progressive global increase in the areal extension of PD epidemic-risk zones (*r*(*t*) *>* 0) in the last decade, irrespective of vineyard distribution (see movies on [57]). This is even more accentuated in the model projections for 2050 which point out a global expansion of PD epidemic-risk zones at different velocities among continents due to climate change (Fig. 5). For example, many important viticulture areas in western Europe included in non-risk or transition zones prior to 1990 are progressively shifting to hotter summers and milder winters and hence would be increasingly suitable for the disease within the extrapolated current scenario. This is further illustrated by a 40% increase of the potential epidemic-risk zone by 2050 with respect to 2020 for Europe and more moderate increases in the United States and in the Southern Hemisphere (Fig. 5). Nonetheless, our model projection for 2050 that includes spatial heterogeneity in the vector distribution, as in Europe, would indicate a lower transmissibility because global change is predicted to have negative effects on *P. spumarius* abundance in Europe [35, 68]. At global scale, there is certain scientific consensus that climate change will follow a general pattern summarised in the paradigm “dry gets drier, wet gets wetter” [69]. In agreement, our model projection for PD on vineyards of Majorca (Spain) suggests shifts to slightly less favourable conditions for Xf_PD_ transmission and an expected progressive decrease in the impact of the disease by 2050. This example and others in Mediterranean islands (see Supplementary Data S4) advocate for certain caution when interpreting climate change projections, especially in other Mediterranean climates of the world, where the complex interactions between humidity and temperature can limit the presence and abundance of vectors (Supplementary Fig. S8).

The scope of our study is constrained by scale limitations, precluding the location-specific complexities surrounding PD ecology. The spatial distribution of the vector is considered only for the *V. vinifera*-Xf_PD_-*P. spumarius* pathosystem in Europe, so the average *R*_0_ numbers could locally differ in other wine-producing regions around the world (Fig. 3), leading to broad local variation in disease incidence where climate is conducive to PD. Such variation is because transmission rates tend to increase exponentially rather than linearly under environmental conditions favouring vector abundance [43], as has been observed at local-scale on vineyards of Majorca [12]. Our study also does not contemplate likely changes within the PD pathosystem. To date, PD is caused by Xf_PD_ (i.e., ST1/ST2), but other genotypes of the subsp. *fastidiosa* or other subspecies and their recombinations could arise in the future with different ecological and virulence traits [19]. On the other hand, new vector species could be accidentally brought in [30], as exemplified with the introduction of the glassy-winged sharpshooter (*Homalodisca vitripennis*) in California, modifying transmission rates and disease incidence in new areas [44]. To capture these uncertainties in relation to the vector, we have performed simulations with *R*_0_ = 8 and *R*_0_ = 16 (Fig. 4). Remarkably, a comparison of PD risk maps for Europe with different *R*_0_ suggests for non-Mediterranean areas the need to stress more surveillance on the introduction of alien vectors rather than in the pathogen itself. This is because, under the current scenario (*R*_0_ = 5) with *P. spumarius* as the main vector, most of the non-Mediterranean vineyards would not support the establishment of PD, but the introduction of new insect vectors with greater transmission efficiency (*R*_0_ = 8) could compensate the climatic layer and increase the risk index above 0. In addition, differences in grapevine varietal response alongside virulence variation among Xf strains may slightly modify PD thermal tolerance limits and therefore locally modulate epidemic intensity (see details in Supplementary Information). Such effect could be seen with cv. Tempranillo, a widely planted variety in northern Spain (Supplementary Table S1); the rate of symptom progress and systemic movement is higher than the average varietal response to Xf_PD_ (i.e., lower *MGDD*), which in addition might imply higher survival rates. This point calls for further testing in the field.

Our model partially explains why PD has not become established in continental Europe and in other main wine-growing regions worldwide during the last 150 years, in contrast to other exotic diseases and pests brought in with native vines from the United States [5– 8]. We suggest that the underlying causes of this low-invasiveness risk in Europe are fundamentally two: (i) a low climatic suitability for chronic PD and (ii) a climatic mismatch between environment conditions suitable for both the vector and the pathogen and their interplay in disease dynamics, similar to the situation recently described for the *V. vinifera*-Xf_PD_-*P. spumarius* pathosystem in northern California [33]. Currently, suitable conditions for the pathogen’s invasion mostly concur in Mediterranean islands and coastlands (Supplementary Data S4). Likewise, similar results would be expected in other Mediterranean climates of main winegrowing regions of the Southern Hemisphere if a vector spatial distribution layer is incorporated in the model simulations (see [57]). Finally, although increasing global warming will extent epidemic-risk zones in all continents, some caution is recommended to not incur in risk overestimation, as we show in the PD risk projections for 2050 in Europe when taking into account the vector spatial distribution; complex interactions between temperature and humidity in the ecology of the vectors may have a great effect in their distribution, abundance and thus transmission capacity [35]. There is an urgent need to fill the knowledge gap on the ecophysiology for each potential vector to downscale PD model predictions to local and regional situations.

## Methods

### Inoculation tests

We carried out an extensive Xf inoculation test on a large number of European vine varieties to evaluate their response to the development of PD symptoms. Fifty-seven rootstock-scion combinations of local, regional and international varieties were pinprick mechanically inoculated with two isolates of Xf. subsp. *fastidiosa* (ST1) recovered from grapevines [25]. Plants were randomly distributed in 12-plant rows along an insect-proof net tunnel exposed to air temperature (Supplementary Table S1). Disease severity was rated by counting the number of symptomatic leaves eight weeks-post-inoculation (wpi) and then every two weeks until the 16th week. Full details on the inoculation conditions, isolates, disease score and statistical analysis are provided in Supplementary Information.

### Modified Growing Degree Days

We generalised the concept of Growing Degree Days (GDD) to account for the specific growth rate of Xf_PD_ as a function of temperature based on the work of Feil & Purcell [38]. A multilinear fit was performed between Xf growth rate and temperature to redefine the classic GDD function into a new metric: the modified growing degree days (MGDD) (see Supplementary Section S2 A). The base, optimal and maximum temperatures defining the MGDD metric are given by *T*_base_ = 12 °C, *T*_opt_ = 28 °C, *T*_max_ = 35 °C and were directly retrieved from Feil & Purcell measures [38]. The mathematical relationship between bacterial population growth and the accumulated MGDD is shown in Supplementary Section S2 B).

### Disease expression associated with temperature

Hourly mean temperature data was recorded over the 3-year trial with an automated weather station (Quimisur, IQ2000) located outside the insect-proof net tunnel. We transformed the cumulative temperatures into units of *MGDD* to characterise the development of PD symptoms [70]. Data of the number of symptomatic leaves and *MGDD* for each plant were pooled across the 36 inoculated grapevine varieties and 57 rootstock-scion combinations to obtain a generalised average pattern of *V. vinifera* response to Xf_PD_. We modelled by survival analysis the likelihood of developing PD symptomatic leaves as a function of *MGDD*, fitting a logistic function. As the event of interest, we assigned the value of five or more symptomatic leaves corresponding to the valley in the bimodal distribution of the number of symptomatic leaves (Supplementary Fig. S1). This valley can be interpreted as the threshold that distinguishes infections that remain stationary and those that accelerate at 12 wpi. A Kaplan-Meier median estimate and the percentiles 10th, 33th, 66th and 90th were used to scale the risk of the total *MGDD* in the parameterised logistic function, ℱ (*MGDD*) (Fig. 1C). In our model ℱ (*MGDD*) represents the cumulative probability of a new infection becoming chronic due to *MGDD* accumulation

### Disease recovery through winter curing

To capture the accumulation nature of the chilling process and differences in climate zones, we determined the global average correlation between *T*_min_ and CDD using 6,487,200 points distributed throughout the planet. We found an exponential relation, *CDD* ∼ 230 exp(− 0.26 *T*_min_), where specifically, *CDD ≳* 306 correspond to *T*_min_ *<* − 1.1 °C. To transform this exponential relationship to a probabilistic function analogous to ℱ (*MGDD*) ranging between 0 and 1, we considered the sigmoidal family of functions 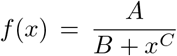 with *A* = 9 *·* 10^6^, *B* = *A* and *C* = 3 (Fig. 1C), fulfilling the limit (*CDD* = 0) = 1, i.e. no winter curing when no cold accumulated, and a conservative 75% of the infected plants recovered at *T*_min_ = − 1.1 °C instead of 100% to reflect uncertainties on the effect of winter curing.

### Global climate data and MGDD/CDD computation

Worldwide temperature data were downloaded from the ERA5-Land dataset [59] with hourly temporal resolution and 0.1^*o*^ spatial resolution using GRIB format. To compute the annual *MGDD* and *CDD* estimates a simple Julia [71] library was built on top of GRIB.jl package [72]. The code can be found in [73]. Data on the vineyard surface in Europe were obtained from the CORINE land-cover map ([74]). For the Northern Hemisphere, the accumulated *MGDD*s were computed from the first of April to the 31st of October, whereas (*CDD*s) were estimated from the first of November to the 31st of March using a 6^*o*^C base temperature [39], and the reverse for the Southern Hemisphere.

### *Philaenus spumarius* species distribution modelling for Europe

SDMs-derived estimation of potential distribution of *P. spumarius* in Europe under current and future (i.e. 2050) climatic conditions were provided by Godefroid et al. [35]. Predictions were obtained using a generalised additive model and two bioclimatic descriptors i.e., a climatic moisture index for the coldest 8-month period of the year and average maximum temperature in spring (March, April and May). Both descriptors reflect physiological constraints acting on life stages of the meadow spittlebug, particularly sensitive to spring temperature and humidity (eggs and nymphs), and were identified as good predictors of *P. spumarius* distribution ([35] and Supplementary Fig. S8).We used the positive relationship found between the climate suitability and abundance of *P. spumarius* adults [35] to assume no climatic constraints on vector population sizes at the optimal climatic conditions *v* = 1. Climatic suitability indices, *v*(*x*), were used to compute a spatially-dependent basic reproduction number, *R*_0_(*x*) = *R*_0_ *v*(*x*). The assumption of the linear dependence between the basic reproduction number and climatic suitability is based on vector-borne epidemic compartmental models (Supplementary Sections S2 F and S4).

### Risk assessment by 2050

Climatic variables were obtained extrapolating the computed *MGDD* and *CDD* time series up to 2050 with annual resolution. The observed trends of the time series were captured using a machine learning based linear regression model while the inter-annual fluctuations where modelled by a Gaussian noise (Supplementary Section S3). Future risk extrapolations were obtained as the average of 10^4^ simulations of this process. A correlative SDM was used to estimate vector spatial distribution in Europe using the global circulation model MIROC5 and greenhouse gas emission scenario rcp45, assuming moderate climate change [35]. Afterwards, the risk was computed following the same simulation procedure previously explained.

## Data accessibility

We provide a library built in Julia to analyse the data outputs of ERA5-Land in GRIB format in [73]. Furthermore, the simulation code and a small reproducible example is provided in [75].

## Acknowledgements

The authors thank the Balearic Islands Official Plant Health Laboratory (LOSVIB) and acknowledge funding from the Spanish Ministry of Science and Innovation through Grants RTI2018-095441-B-C22 (SuMaEco) (AGR and MAM) and RTI2018-093732-B-C22 (PACSS) (JJR) funded by MCIN/AEI/10.13039/501100011033 and by ERDF “A way of making Europe”, MDM-2017-0711 (AGR, JG, JJR and MAM) funded by MCIN/AEI/10.13039/501100011033; The Ministry of Agriculture, Fishery and Food of the Government of the Balearic Islands under Grant (MEPRO 11876/2019). MG received the grant Ayudas a la Atracción de Talento Investigador (Ref: 2018-T2/ BIO-1137) funded by la Comunidad de Madrid.

## Author contributions

AGR, AF, JB, JJR, MAM and EM planned and designed research. MM and EM carried out the inoculation experiments. AGR, JG, JJR and MAM conceptualised the mathematical model. AF and MG developed ecological models for the distribution of *Philaneus spumarius*. AGR, JG, JB and MG built the risk maps. AGR and EM wrote the original draft. AGR, MAM, JJR and EM reviewed and edited the final manuscript.

