## Supplementary Material for "Global predictions for the risk of establishment of Pierce’s disease of grapevines"

**This PDF file includes:**

- Supplementary text
- Figures S1 to S11
- Tables S1 to S9
- SI References

### CONTENTS

List of Figures 3

List of Tables 4

S1. Inoculation tests on European grapevine varieties 5

S2. Modelling climate suitability for PD 6

Figures 12

Tables 23

References 32

### LIST OF FIGURES

S1 **Factors influencing *Xf-Philaenus spumarius-Vitis vinifera* pathosystem.(a)** Virulence differences between *Xf* subsp. *fastidiosa* isolates on grapevines. Bars represent the mean number of symptomatic infected leaves four months after inoculating. Both isolates XYL2055/17 ($n = 316$ inoculated plants) and XYL2177/18 ($n = 260$) were collected from vineyards on Majorca. Scores were pooled among the 21 varieties inoculated;**(b)**conceptual graph of the population dynamics of *P. spumarius* on vineyards in Majorca and the effect on winter curing. Blue: density function of *P. spumarius*; red line: proportion of *P. spumarius* carrying Xf; blue line: proportion of plants recovering according the time they are infected;**(c)** bimodal density function of the number of symptomatic leaves. Blue dash line marks 5 symptomatic leaves; and **(d)**correlation between the upward and downward movement of $Xf_{PD}$ within the canes from the inoculation point. Each point depicts mean distance travelled in both directions by the bacteria. . . . . . 12

S2 **Experimental setup.** Greenhouse facilities and general view of its interior and the arrangement of the vine plants. The metallic structure is covered with an anti-thrips mesh. . . . . . 13

S3 **Relationship between MGDD and temperature.** Contribution to the $MGDD$ resulting from the fitting to the data in (1). The original Arrhenius plot, $\log k$ *vs.* $1/T$ in kelvin was converted to a linear dependence in Celsius temperature $t$ (cf. Eq. (S3)) . . . . . . 14

S4 **Trends in the risk-epidemic zones during the 1981-2019 period (A) and the areas encompassed below the $CDD < 314$ line (B) comprising land areas between $103^o$W and $70^o$W of the United States** . . . . . . 15

S5 **ROC curve illustrating the model validation procedure with spatiotemporal data from PD distribution in the US.** TPR is the *true positive rate* and FPR the *false positive rate.* The model accuracy reaches its optimum in $R_0 \approx 8$ by maximising the true positive rate and minimising the false positive rate. The spatiotemporal PD distribution in the US was obtained from data collected from publications between 2001 and 2015. . . . . . 16

S6 **Fitting a $SIR$ model to the progress of the almond leaf scorch disease in the Balearic Islands from 1993 onward**.The best match was obtained with $R_0 = 11.2$. Points represent an estimate of the proportion of infected trees (incidence) from dendrochronological analysis and detection of Xf DNA in growth rings by qPCR. The incidence of ALSD in 2012 and 2017 was independently validated by field and Google Map Street View image observations. . . . . . 17

S7 **Model validation for an $R_0 = 8$ scenario with presence/absence data (black/white stars) of PD in the United States.** Panel (A) corresponds to data from California in 2015 while the other panels show data from 2002, 2005, 2006 and 2001 (respectively) in the east of the United States. The last panel clarifies the validation zones previously mentioned. . . . . . 18

S8 **Average climatic suitability for *Philaenus spumarius* in Europe.** The map shows the climatic suitability of the vector estimated from a generalized additive model of insect distribution and the correlation of two bioclimatic descriptors, a climatic humidity index for the period of 8 coldest months of the year and the average maximum temperature in spring. . . . . . 19

S9 **Trends in $MGDD$ (A) and $CDD$ (B) values and oscillations during 1981-2019 for seven wine regions with different climates from Europe and the US.** $MGDD$s show a slight upward trend and lesser oscillations than the $CDD$s. . . . . . 20

S10 **Determination of $MGDD$ and $CDD$ metric trends and future projections for two different European regions.** The $MGDD$ (A) and $CDD$ (B) trends show steeper slopes in the temperate climate of Bordeaux than in the Mediterranean climate of Lecce. . . . . . 21

S11 **Interannual climatic variability extrapolations of $MGDD$ (A) and $CDD$ (B) for Bordeaux.** A linear model was fitted using Sklearn's LinearRegression module in Python and the interannual climatic variability was included as a Gaussian noise distribution by calculating the mean and fluctuations of the variance around $MGDD$ and $CDD$ trends. . . . . . 22

### LIST OF TABLES

S1 **Summary of the inoculation tests on grapevine varieties ranked from most to less susceptible in the disease index.** Thirty-six local, regional and international varieties were screened in combination with eight rootstocks. The number of symptomatic leaves were counted 16 weeks after inoculation and infections confirmed by qPCR. DI: disease index; AUDCP: area under the disease progress curve. . . . . . 23

S2 **PD risk areas in Europe after running the model under a $R_0 = 5$ scenario and a homogeneous spatial vector distribution.** The epidemic-risk zones are classified according to the relative disease growth rates defined by the risk index, as very low, low, moderate and high growth rates. The total risk refers to the sum of the epidemic-risk zones . . . . . . 25

S3 **PD risk areas in the United States after running the model under a $R_0 = 8$ scenario and using a homogeneous spatial vector distribution.** The epidemic-risk zones are classified according to the relative disease growth rates defined by the risk index, as very low, low, moderate and high growth rates. The total risk refers to the sum of the epidemic-risk zones . . . . . . 26

S4 **Potential distribution of PD in other world winegrowing regions.** In most areas of China and Australia *Vitis vinifera* is not cultivated and epidemic-risk zones with high growth rate correspond mainly to tropical areas in China, Australia, South Africa and Argentina . . . . . . 27

S5 **Predicted PD risk areas for the US in 2050 considering a $R_0 = 8$ scenario and a homogeneous spatial vector distribution.**The epidemic-risk zones are classified according to the relative disease growth rates defined by the risk index, as very low, low, moderate and high growth rates. The total risk refers to the sum of the epidemic-risk zones . . . . . . 28

S6 **Predicted PD risk areas in Europe in 2050 after running the model under a $R_0 = 5$ scenario and a homogeneous spatial vector distribution.** The epidemic-risk zones are classified according to the relative disease growth rates defined by the risk index, as very low, low, moderate and high growth rates. The total risk refers to the sum of the epidemic-risk zones . . . . . . 29

S7 **PD risk areas in Europe PD risk areas in Europe after running the model under a $R_0 = 5$ scenario and a spatial heterogeneous vector distribution (climatic suitability).** The epidemic-risk zones are classified according to the relative disease growth rates defined by the risk index, as very low, low, moderate and high growth rates. The total risk refers to the sum of the epidemic-risk zones . . . . . . 30

S8 **Surface of European vineyards in risk of PD given by the intersection of (Corine-Land-Cover) and the projected model in the ERA5-land data under a $R_0 = 5$ scenario with the layer of vector climatic suitability.**The epidemic-risk zones are classified according to the relative disease growth rates defined by the risk index, as very low (0.1-0.33), low (0.33-0.66), moderate (0.66-0.9) and high exponential growth rates ($> 90$). The total risk refers to the sum of the epidemic-risk zones. . . . . . . 31

S9 **Predicted PD risk in 2050 in European vineyards (Corine-Land-Cover) considering a $R_0 = 5$ scenario and the vector climatic suitability.**The epidemic-risk zones are classified according to the relative disease growth rates defined by the risk index, as very low, low, moderate and high growth rates. The total risk refers to the sum of the epidemic-risk zones . . . . . . 32

### S1. INOCULATION TESTS ON EUROPEAN GRAPEVINE VARIETIES

**Plants**. Grapevine saplings were annually supplied from a nursery of mainland Spain (Viveros Villanueva Vides, SL), consisting of one-year-old rootstocks grafted in winter with dormant grapevine cultivars, and grown in 20-L plastic pots with a standard potting mix. Fifty-seven rootstock-scion cultivar combinations were used in the inoculation assay (Table S1). Potted plants were randomly distributed in 12-plant rows along an insect-proof tunnel exposed to air temperature and daily dip-irrigated to field capacity, fortnightly sprinkled with a slow-release fertiliser and treated with insecticides and fungicides when needed until the end of the experiment. Two weeks before the onset of the inoculation assay, leaf samples of all plants were collected and tested for the presence of Xf through qPCR as described elsewhere [1].

**Isolates and inoculation**. We used for the inoculation experiment two isolates of Xf. subsp. *fastidiosa* (ST1) recovered from grapevines: XLY 2055/17 (GenBank WGS: QTJS01) and XYL2177/18 (JAAGVM01) [2, 3]. In the third-year assay we included an isolate of Xf subsp. *multiplex* ST81 XYL1981/18 (JAAGVE01) to test whether other strains in Majorca could cause PD as well. Isolates were grown on BYCE medium at 28$^{o}$C for 7-10 days, following EPPO protocols [4]. Cells were collected by scraping the colonies and suspending them in a1.5 ml Eppendorf tubes each with 1 ml of phosphate-buffered saline (PBS) solution until obtaining a turbid ($10^8 - 10^9$ cell/ml) suspension. Plants were mechanically inoculated by pin-prick inoculation [5] with slight modifications. A 10-$\mu l$ drop of the bacterial suspension was pipetted on the leaf axil and punctured five times with an entomological needle. Eight-nine replicates per scion-rootstock combination were inoculated with the bacterial suspension and four-three plants per cultivar with a drop of PBS as a control at the end of May. Inoculation was repeated two weeks thereafter by piercing the next leaf axil above that previously inoculated [1].

**Disease score.** Disease severity was rated by counting the number of symptomatic leaves eight weeks post-inoculation (WPI) and then biweekly until the 16th week. A disease index was calculated according to Su et al. [6]. To determine the basipetal and systemic movement of $\mathrm{Xf_{PD}}$, we counted the number of symptomatic leaves below the point of inoculation from the same stems or any stem below at 12 WPI. Symptomatic and asymptomatic plants were tested by qPCR for Xf infection at 12 WPI taking the petiole of the second and fifth leaf above the point of inoculation. On the 14th week, five leaves per plant of all inoculated plants were used for $\mathrm{Xf_{PD}}$ isolation, as described below. Those plants for which the qPCR was negative and $\mathrm{Xf_{PD}}$ could not be isolated were treated as not infected.

**Data analysis**. All statistical analyses on disease scores were carried out using R. 3.5.2 version software [7]. We used the functions glm and glmer in the R package lme4 [8] for fitting Generalised Linear Models and Generalised Linear Mixed Models (GLMMs) in the analysis of disease incidence and severity in the inoculation assays. In all tests, we modelled the response variable (i) disease incidence with the binomial error (logit-link function) and (ii) disease severity with the Poisson error (log-link function). A within-subject (repeated measures) factorial design was performed to evaluate differences in disease severity over time among different cultivar-rootstock combinations. Cultivars-rootstock and time were treated as fixed factors and plant subjects as a random effect. Rootstock and time were analysed as fixed factors and plant subjects as a random effect. Controls were excluded from the analysis, as lesions did not develop on them. For each cultivar, we calculated the area under the disease progress curve (AUDUP) from weeks-post-inoculation and disease index using the package Agricolae. To test whether genotypes within the ST1-grapevine population vary in virulence, we included a second strain (XYL2177/18) in the 2019 inoculation experiment.

**Varietal response to Xf**. In our three-year inoculation tests, we included a representative number of local and international varieties (Table S1). In total, among 886 inoculated-grapevine plants comprising 36 varieties in 57-unique combinations (scion-rootstock), 86.1% ($n = 764$) of them developed typical PD symptoms at 16 WPI. In contrast, none of the grapevine plants inoculated with the strain XYL 1981/17, ST81 subsp. *multiplex* presented symptoms. The results of the pathogenicity tests on European grape varieties are shown in Table S1.

Overall, European *V. vinifera* varieties exhibited significant differences in their susceptibility to $\mathrm{Xf_{PD}}$, which could imply differences in risk of PD establishment at regional scale (Table S1). When compared between grape major phenotypic groups, red grape varieties were 1.45 times more prone to $\mathrm{Xf_{PD}}$ infection than white grape varieties ($\chi^2 = 41.58$, $df = 1$, $P = 1.072 \times 10^{-10}$), while symptoms were 36.7% more severe in red grapes than in white grape cultivars ($\chi^2 = 554.54$, $df = 1$, $P = 2.2 \times 10^{-16}$). In addition, we probed whether $\mathrm{Xf_{PD}}$ strains isolated from grapevines in Majorca differ in their virulence pooled across all grapevine varieties, finding significant differences in virulence ($\chi^2 = 68.73$, $df = 1$, $P = 2.2 \times 10^{-16}$) and infectiveness ($\chi^2 = 8.07$, $df = 1$, $P = 0.0045$) (Fig. S1).

Early-season Xf-infections on grapevines are considered to be more likely to survive the following year than late-season infections [9, 10]. By contrast, varieties developing symptoms later on may affect pathogen acquisition efficiency by vectors and thus decrease the rate of disease transmission. We found a positive correlation ($F_{1,28} = 39.58$,

$P < 0.001$; $R^2 = 0.57$) between the number of symptomatic leaves formed above the point of inoculation and those formed below (Fig. S1). This acropetal/basipetal ratio of infected leaves is indicative of systemic movement of the pathogen and of a greater probability that infections on vines showing a lower number of symptomatic leaves will be more likely eliminated by winter pruning or by low temperatures [11]. As a result, we assumed in our model that Xf-infected plants that develop fewer symptomatic leaves at the end of 16 weeks of incubation will contribute less to the spread of the disease within vineyards (Fig. S1).

### S2. MODELLING CLIMATE SUITABILITY FOR PD

#### A. Modified Growing Degree Days (MGDD) from Arrhenius Equation

Feil and Purcell estimated Xf growth rate as a function of temperature, $\sigma(T)$, using an Arrhenius' Law, i.e. $\ln K \sim -1/T$ (see Fig. 3 in [9]). The overall dependence on $T$ is nonmonotonic with two different types of behaviour: i) $k$ grows with $T$ until a maximum value is attained at $T = 28\,^{\circ}\mathrm{C} = 301.15\,\mathrm{K}$; and ii) $k$ decreases beyond the maximum. Growth is zero beyond the lowest and highest threshold temperatures.

The mathematical form of the Arrhenius' Law dependence between the growth rate $k$ and the absolute temperature $T$ reads as follows,

$$k = A\exp(-E/T)\ , \tag{S1}$$

where $A$ is a pre-exponential factor and $E$ an activation energy in units of the Boltzmann constant $k_B$. The original use of this equation is for the rate constant of a chemical reaction that increases monotonically with $T$, and so $E > 0$. To fit the non-monotonic whole growth behaviour of Xf, we considered two Arrhenius functions with opposite signs in the activation rate,

$$k = A_1\exp(-E_1/T) + A_2\exp(+E_2/T)\ , \tag{S2}$$

where $E_1 > 0$ and $E_2 < 0$.

Now let us denote by $t$, the temperature in Celsius, $t = T - 273.15$. Importantly, within the bacterial temperature growth range (10-36 °C) in the Arrhenius equation Eq. (S2), $t$ is quite small respect to $b = 273.15$, the absolute (Kelvin) temperature corresponding to 0 °C in $T = 273.15 + t$. The two exponents in Eq. (S2) can be approximated now as,

$$\begin{aligned} k = A\exp\left(-\frac{E}{b+t}\right) = A\exp\left(-\frac{E}{b(1+t/b)}\right) \approx A\exp\left[-\frac{E}{b}\left(1-\frac{t}{b}\right)\right] = A\exp\left(-\frac{E}{b}\right)\exp\left(\frac{E}{b^2}t\right) \approx \\ A\exp\left[-\frac{E}{b}\right]\left(1+\frac{E}{b^2}t\right) = A\exp\left(-\frac{E}{b}\right) + A\frac{E}{b^2}\exp\left(-\frac{E}{b}\right)t = B + Ct\ , \end{aligned} \tag{S3}$$

where we assume that $t/b = t/273.15 \ll 1$ and $(Et/(b^2) = (Et)/(273.152) \ll 1$, whereas $B$ and $C$ are constants. In particular, $C > 0$ if $E > 0$ fits the region before the maximum in which the growth rate increases, while $C < 0$ if $E < 0$ fits the region after the maximum where $k$ decreases. The positive/negative sign stems from the the coefficient of the linear term in $t$, $E/b^2$.

Each exponential in Eq. (S2) can be expressed with a simple straight line, valid in our temperature range. This approach can be extended by adding more exponential terms in Eq. (S2) to further improve the fit with a multi-linear dependence between $\mathrm{Xf}_{\mathrm{PD}}$ growth rate and temperature, obtaining a function proportional to $\mathrm{Xf}_{\mathrm{PD}}$ growth rate $F(T) = C\cdot\sigma(T)$ (see Fig. S3).

Now, this multi-linear fit to the $\mathrm{Xf}_{\mathrm{PD}}$ growth rate can be used to redefine the classical Growing Degree-Days (GDD) metric into the new Modified Growing Degree Days (MGDD). *GDD*s are computed as the integral of a particular function of temperature

$$GDD(t) = \int_{t_0}^{t} f(T(t))\mathrm{d}t \tag{S4}$$

where $f(T)$ is defined as

$$f(T(t)) = \begin{cases} T(t) - T_{\mathrm{base}} & \text{if } T \geq T_{\mathrm{base}} \\ 0 & \text{if } T < T_{\mathrm{base}} \end{cases} \tag{S5}$$

Considering different slopes relating $Xf_{PD}$ growth rate and temperature at different temperature intervals, as shown in Fig. S3, we modified this particular function to now account for $Xf_{PD}$ growth,

$$MGDD(t) = \int_{t_0}^{t} F(T(t))\mathrm{d}t = C \cdot \int_{t_0}^{t} \sigma(T(t))\mathrm{d}t \tag{S6}$$

We wish to stress that the use of a multilinear form to represent MGDDs Fig. S3 stems from the fundamental temperature dependence of the kinetics of bacterial growth as described by Arrhenius equation, and is not an arbitrary simplified representation. Moreover, the MGDD function is fitted using the whole set of data published in [9], and is not simply based on the knowledge of the cardinal temperatures, as customarily done when writing smooth interpolating functions with the sole input of the cardinal temperatures (see, e.g., [12]).

### B. Relation between MGDD and within-plant bacterial population

The usual growth cycle of bacteria consists of several phases (lag, exponential, stationary and death phase), being of most interest to environmental microbiologists the interval between the lag and the onset of the stationary phase [13]. During the exponential phase, the rate of increase of cells is proportional to the number of cells present at any particular time. Thus, the evolution of the bacterial population, $N$, over time is given by the following differential equation,

$$\frac{\mathrm{d}N}{\mathrm{d}t} = \sigma N \Longrightarrow N(t) = N_0 \cdot \exp(\sigma t) \ , \tag{S7}$$

where $\sigma$ is the specific growth rate constant.

As shown in the previous section, the growth rate of Xf has specific temperature dependence, $\sigma(T)$. In our study, temperature varies over time, so we can write the growth rate as a time-dependent quantity, $\sigma(T(t))$. With this, the evolution of the bacterial population will be given by

$$N(t) = N_0 \exp\left(\int_{t_0}^{t_f} \sigma(T(t))\mathrm{d}t\right) \ . \tag{S8}$$

Recalling Eq. (S6) we can write the previous equation as

$$N(t) = \frac{N_0}{C} \cdot \exp(MGDD(t)) = C' \cdot N_0 \exp(MGDD(t)) \tag{S9}$$

Indeed, the same can be done considering the logistic differential equation (that includes the stationary phase),

$$\frac{\mathrm{d}N}{\mathrm{d}t} = \sigma(T(t)) \cdot N \cdot \left(1 - \frac{K}{N}\right) \tag{S10}$$

whose solution is

$$N(t) = \frac{K}{1 + \exp\left(\int_{t_0}^{t_f} \sigma(T(t))\right)} \tag{S11}$$

and using Eq. (S6) it can be rewritten as

$$N(t) = \frac{K}{1 + C\exp(-MGDD(t))} \tag{S12}$$

and thereby the bacterial population after a given time $t$ is related to the $MGDD$ by Eq. (S12).

**Note:** We are assuming a correspondence between *in vitro* and *in planta* growth rates of Xf.

#### C. Epidemiological and theoretical basis

A standard SIR model was considered as a basis to asses the risk of PD outbreaks worldwide (see Section S4 for an analytical derivation of the relation between a vector-borne disease model and a standard SIR model). The model is represented by the following three equations,

$$\begin{aligned}\dot{S} &= -\beta SI/N \\ \dot{I} &= \beta SI/N - \gamma I \\ \dot{R} &= \gamma I\ ,\end{aligned} \tag{S13}$$

where $S$ is the susceptible host population, $I$ the infected population, $R$ the dead population and the total population $N$ is conserved, $S + I + R = N$, as hosts die only when they contract the disease. The transmission of the disease from infected hosts to susceptible ones is mediated by the *transmission rate* $\beta$ while the death of infected individuals is regulated by the *mortality rate* $\gamma$.

Analysing the non-trivial fixed point, $\boldsymbol{x} = (N, 0, 0)$, it can be proved the existence of an epidemic threshold. As $S$ is a monotonically decreasing function, which implies $S(t) < S_0$, one can write the following relation,

$$\frac{\mathrm{d}I}{\mathrm{d}t} = I\left(\beta S/N - \gamma\right) \leq I\left(\beta N/N - \gamma\right) = \gamma I\left(\beta/\gamma - 1\right) = \gamma I\left(R_0 - 1\right)\ , \tag{S14}$$

where $R_0 = \beta/\gamma$. Thus, $R_0 < 1$ implies $dI/dt < 0\ \forall t$ and $I_0 > I(t)$ as $t \to \infty$, basically meaning that the epidemic dies out, while for $R_0 > 1$, $I(t)$ grows initially until $S(t_c) = \gamma/\beta$, at which $\dot{I}(t_c) = 0$, and the epidemic starts waning out. $R_0$ corresponds to the so-called *basic reproduction number* and measures the number of secondary infections given by a primary infection in a fully susceptible population.

We wish to model the risk of PD establishment in a susceptible (healthy) population. For this, we characterised the maximum growth rate of the epidemic, when $S(t) \sim S(0)$. Thus, the growth is well approximated under these conditions with the (linearised) differential equation,

$$dI/dt = \beta SI - \gamma I \approx \gamma I(\beta N/\gamma - 1) = \gamma I(R_0 - 1)\ . \tag{S15}$$

where we have assumed the initial conditions, $S_0 \approx N$, $I(0) \approx 0$ and $R(0) = 0$. This linear differential equation can be integrated exactly,

$$I(t) = I(0)\exp(\gamma(R_0 - 1)t)\ . \tag{S16}$$

As explained in the main text, to account for the effect of temperature in the epidemic process we modify the previous expression as follows

$$I(t) = I(0)\exp(\gamma(R_0 - 1)t) \cdot \mathcal{F}\left(MGDD(t)\right) \cdot \mathcal{G}\left((CDD(t)\right) = I(0)\exp(\gamma(R_0 - 1)t) \cdot \Pi(t)\ , \tag{S17}$$

where $\Pi(t) = \mathcal{F}(MGDD(t)) \cdot \mathcal{G}(CDD(t))$ is the cumulative probability of chronic infection that depends on temperature.

#### D. Determination of $R_0$ for Europe

Unlike the validation of our model based on the distribution of PD in the USA, there are no spatiotemporal data on PD outbreaks available in Europe to estimate $R_0$. One way to solve this problem is to use data on the incidence of almond leaf scorch disease in Majorca to fit a SIR model and obtain an approximation of $R_0$. The initial date of introduction and progression of the almond leaf scorch epidemics is well characterised and both diseases are transmitted by *P. spumarius* [1, 3]. Using $\gamma = 1/14\text{years}^{-1}$ as the mortality rate [3], the best fit was provided by $\beta_{\text{opt}} = 0.8$, giving rise to $R_0 = 11.2$, which is in good agreement with the order of magnitude of $R_0 = 8$ in the United States (Fig. S5). To find a proper scenario for PD in Europe, we considered a constant transmission rate $\beta_{\text{opt}}$[1] and applied an average mortality rate $\gamma \sim 1/5\text{years}^{-1}$ of PD infected vines [14], which gives rise to $R_0 = 4$. Finally, we used the information of the climate suitability for the vector in Majorca ($\approx 0.8$ on average) to determine a baseline scenario for Europe, $R_0 = 4/0.8 = 5$. Thus we can argue that $R_0 = 5$ is a good proxy for modelling the establishment of PD in Europe. The use of $R_0 = 5$ is furthermore corroborated by the reasonability of the predictions obtained.

---

[1] As the vector that transmits both ALSD and PD is the same (*Philaenus spumarius*) we considered that the transmission rate should not vary much.

#### E. Simulation details

To assess the risk of *Xf* establishment in vineyards, we performed spatiotemporal simulations for the world's largest wine-growing areas. The cell size of the abstract grid was determined by the resolution of the data collected from ERA5-Land, 0.1 °×0.1 °, so the spatial resolution is approximately 9 km in the latitudes of the Mediterranean basin. A small initial infected-plant population was introduced annually into each cell assuming that if the conditions are or become favourable the disease will propagate locally. We chose $I_i(0) = 1$ to re-scale the results to any initial population size, and implemented Eq. (S17) in each cell. Simulation time was discretised in years and computed in two steps incorporating summer ($F(MGDD)$) and winter ($F'(CDD)$) periods. To implement Eq. (S17) we took into account that the $MGDD$ and $CDD$ differ at each time step; thereby it required to convert Eq. (S16) into a mathematical map. The equation can be expressed as,

$$I(t) = I(0)\exp(\gamma(R_0 - 1)t) = I(0)\left[\exp(\gamma(R_0 - 1))\right]^t \ , \tag{S18}$$

where $t$ is the discrete time in years, so that

$$I(t-1) = I(0)\left[\exp(\gamma(R_0 - 1))\right]^{t-1} \ , \tag{S19}$$

and, thus,

$$I(t) = I(t-1)\exp(\gamma(R_0 - 1)) \ . \tag{S20}$$

The discretized form of Eq. (S17) is then

$$I(t_i) = I(t_{i-1})\exp(\gamma(R_0 - 1)) \cdot F(MGDD(t_i)) \cdot F'(CDD(t_i)) \ . \tag{S21}$$

A risk index was created to represent the relative velocity of PD local exponential propagation,

$$r(\tau) = \max\left\{\frac{\log(I(\tau)/I(0))}{\gamma(R_0 - 1)\tau}, -1\right\} \ , \tag{S22}$$

where $\tau$ is the simulated time, $R_0$ is the basic reproduction number and $I(0)$ the initial condition (initial number of infected plants). The index ranges from -1 to 1 as the maximum risk value always occur under optimal climatic conditions ($F(MGDD) = F(CDD) = 1$) and thus $I(\tau) = I(0)\exp((R_0 - 1)\tau)$. The minimum risk was intentionally cut off in -1 to use a symmetric scale, as otherwise the logarithmic scale is unbounded.

The numerator of the risk index defined in Eq. (S22) is formally similar to the definition of Lyapunov exponents (LEs), which characterise predictability in chaotic systems (the denominator normalises this quantity to its maximum value). This is not surprising because both the risk of Eq. (S22) and the growth of perturbations in chaotic systems correspond to an exponential process. Following this analogy, we would expect a growing exponential process in the risk of establishment if $r > 0$, while a decreasing exponential that goes to 0 would denote no risk if $r < 0$. However, Lyapunov exponents are (normally) calculated for autonomous (i.e. unforced, and so *steady*) dynamical systems, while Eq. (S21) has 2 forcing terms (i.e. is non-autonomous). The result is a non-exponential behaviour found when $|r|$ is small. So beyond the expected regions with growing exponential and decreasing exponential behaviour, we find a transition zone, where the system is oscillatory and not exponential, as neither growth in more auspicious years for $\mathrm{Xf}_{\mathrm{PD}}$ or decrease in less auspicious ones prevails, and neither of the growing or decreasing pure exponential behaviours manifests.

We define the borderlines of this transition region by $I(\tau) \le 10 \cdot I(0)$ in the southern boundary and $I(\tau) \ge 0.05 \cdot I(0)$ in the northern one when $\tau = 40$ years. Basically, for the upper boundary we assume that if an initial infection is multiplied by 10 after 40 years, then the exponential growth would be unstoppable. Conversely, if an initial introduction of infected individuals decays more than a 95% of its original value after 40 years, we then assume that the exponential decay would continue and clearly PD cannot be established. Since $\tau, \gamma$ are fixed, the limits of the transition zones depend on $R_0$ and it is given by the risk index instead of the number of infected plants as follows,

$$\begin{aligned} \text{Upper limit:} \ r_{\text{trans}}^{\max} &= \frac{\log(10)}{\gamma\,(R_0 - 1)\,\tau} \\ \text{Lower limit:} \ r_{\text{trans}}^{\min} &= \frac{\log(0.05)}{\gamma\,(R_0 - 1)\,\tau} \end{aligned} \ . \tag{S23}$$

For instance, with $\gamma = 0.2\,\text{years}^{-1}$, $\tau = 39\,\text{years}$ and $R_0 = 5$ (values used for Europe) the transition zones are delimited by $-0.09 < r(\tau) < 0.075$. So, the model outputs can be associated to the following behaviours:

1. Epidemic-risk zones: $r(\tau) > r_{\text{trans}}^{\text{max}}$. The risk index $r_j(\tau)$ is ranked as high ($r_j(\tau) > 0.9$), moderate (0.9-0.66), low (0.66-0.33) and very low (0.33-$r_{\text{trans}}^{\text{max}}$).
2. Transition-risk zones: $r_{\text{trans}}^{\text{min}} < r(\tau) < r_{\text{trans}}^{\text{max}}$. In this zone the incidence, $I(t)$, predicted by the model does not grow clearly, but neither it does disappear, and incidence oscillates. This region is expected to be very sensitive to changes induced by climate change, and transit to epidemic-risk zones with low growth rate.
3. Non-risk zone: $r(\tau) < r_{\text{trans}}^{\text{min}}$. Incidence decrease exponentially due to the combined effect of the $MGDD$ and $CDD$, or to the (low) vector abundance in the case of predictions for Europe. Cells in this region with $r_j$ not far from $-0.1$ could become transitional due to the effect of climate change.

#### F. Vector distribution influence

Information on the climatic suitability of the vector *P. spumarius* [15] was used to modulate the value of the basic reproduction number. We assumed a linear dependence of $\beta$, the transmission rate, with the vector climatic suitability resulting in each of the model cells,

$$R_0(x) = \frac{\beta v(x)}{\gamma} = R_0 \cdot v(x) \ , \tag{S24}$$

where $x$ illustrates the space dependence.

In Section S4 we show an analytical derivation of the linear dependence between $R_0$ and the vector population (i.e. the number of vectors). Then, assuming that climatic suitability (i.e. probability of presence) is directly related to the number of vectors we obtain the linear scaling between $R_0$ and climatic suitability for vectors.

### S3. FUTURE RISK EXTRAPOLATION

To project PD risk in a climate change scenario, historical $CDD$ and $MGDD$ data were calculated to generate annual time series for each location recorded in the data set. To obtain the time trend of the variables in each pixel, a linear model was fitted using Sklearn's LinearRegression module in Python [16]. The interannual climatic variability was also included as a Gaussian noise distribution by calculating the mean and fluctuations of the variance around the trend of the $MGDD$ and $CDD$ metrics for any record in the data set. We show in Fig. S10 the determination of the trend of the metrics $MGDD$ and $CDD$ for Lecce and Bordeaux. Fig. S11 shows three realisations to extrapolate the $MGDD$ and $CDD$ metrics for Bordeaux after applying Gaussian noise to the trend. This risk extrapolation to 2050 implies a linear extrapolation of past MGDD and CDD tendencies. Note that because $MGDD$ and $CDD$ functions are nonlinear this is just a rough approximation to the future risk, as non-linearities could play a major role in a climate change scenario.

### S4. SIR MODEL AND $R_0$ LINEAR SCALING WITH VECTOR POPULATION FROM A VECTOR-BORNE DISEASE MODEL

We show how a linear scaling between vector population and the basic reproduction number can be obtained from a vector-borne disease model. Moreover, a SIR model can be derived from the same vector-borne disease model (under some assumptions).

In a model defined according to the following processes,

$$S_H + I_V \xrightarrow{\beta} I_H + I_V \quad I_H \xrightarrow{\gamma} R_H \quad S_V + I_H \xrightarrow{\alpha} I_V + I_H \quad S_V \xrightarrow{\mu} \varnothing \quad I_V \xrightarrow{\mu} \varnothing \ , \tag{S25}$$

where the birth of new susceptible vectors is described as a source term, the host-vector compartmental model can be written as,

$$\begin{aligned} \dot{S}_H &= -\beta S_H I_v / N_H \\ \dot{I}_H &= \beta S_H I_v / N_H - \gamma I_H \\ \dot{R}_H &= \gamma I_H \\ \dot{S}_v &= \delta C - \alpha S_v I_H / N_H - \mu S_v \\ \dot{I}_v &= \alpha S_v I_H / N_H - \mu I_v \ , \end{aligned} \tag{S26}$$

when a standard incidence [17] is considered.

The model describes infection of susceptible hosts ($S_H$) at a rate $\beta$ through their interaction with infected vectors ($I_v$), while susceptible vectors ($S_v$) are infected at a rate $\alpha$ through their interaction with infected hosts ($I_H$). Infected hosts exit the infected compartment at rate $\gamma$, while infected vectors stay infected the rest of their lifetime, since they are not affected by the pathogen.The model assumes that vectors die naturally (or disappear from the population by some mechanism) at rate $\mu$ and are born (appear) at a constant rate $\delta$, being susceptible. The constant term $C$ sets the scale of the stationary value of the vector population.

#### A. Linear scaling of $R_0$ with vector population

The standard methods of calculation of $R_0$ are based in the linear stability analysis of the disease-free equilibrium, either directly, through the linear analysis of the fixed point that yields the stability condition from which $R_0$ can be obtained, or using the Next Generation Method (NGM) [18] that provides directly $R_0$ by solving a suitable linear problem. The disease free equilibrium of the model (the fixed point) is given by $I_H = I_v = 0$ yielding $\dot{S}_v = 0 \Longrightarrow S_v = \delta C/\mu = N_v^*$, where $N_v^*$ is the stationary value of the vector population.

As shown in [19], both methods yield the following relation for the basic reproduction number,

$$R_0 = \frac{\beta\alpha}{\gamma\mu}\frac{C}{N_H}\frac{\delta}{\mu}\frac{S_H(0)}{N_H} = \frac{\beta\alpha}{\gamma\mu}\frac{N_v^*}{N_H}\frac{S_H(0)}{N_H} \ , \tag{S27}$$

in which the basic reproduction number scales linearly with the vector population.

#### B. Reduction to a SIR model

In a time-scale where the vector population changes faster than the host population (a good approximation for $Xf_{\mathrm{PD}}$-related diseases), the former will almost instantaneously reach the stationary value. Thus, if $1/\mu \ll 1/\gamma$, or equivalently if $\mu \gg \gamma$, we can rewrite the time derivative of the vector infected population as

$$\epsilon \dot{I}_v = \frac{\alpha}{\mu} S_v \frac{I_H}{N_H} - I_v \ , \tag{S28}$$

with $\epsilon = 1/\mu$ being a small parameter. Then, $\dot{I}_v$ can be neglected and the infected vector population can be obtained from the relationship,

$$I_v \approx \frac{\alpha}{\mu}\frac{S_v I_H}{N_H} \ . \tag{S29}$$

Furthermore, if $\lambda N_H \gg I_H$ (which is indeed plausible in this limit) the model can be written as a SIR model with constant coefficients,

$$\begin{aligned} \dot{S}_H &= -\beta_{eff}\frac{S_H I_H}{N_H} \\ \dot{I}_H &= \beta_{eff}\frac{S_H I_H}{N_H} - \gamma I_H \\ \dot{R}_H &= \gamma I_H \ , \end{aligned} \tag{S30}$$

where $\beta_{eff} = \dfrac{\beta'}{\lambda} = \dfrac{\beta\alpha N_v^*}{\mu N_H}$.

Note that in the SIR model reduction, the effective $\beta_{eff}$ coefficient depends linearly on the vector population $N_v^*$.

**FIGURES**

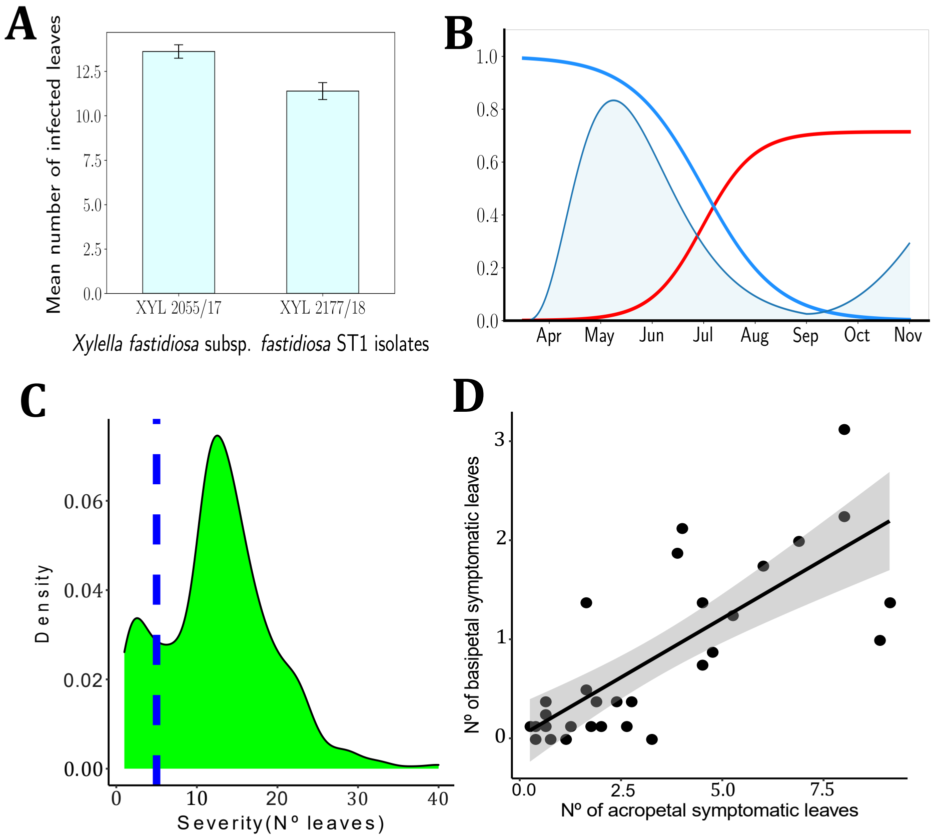

FIG. S1. **Factors influencing *Xf-Philaenus spumarius-Vitis vinifera* pathosystem.(a)** Virulence differences between *Xf* subsp. *fastidiosa* isolates on grapevines. Bars represent the mean number of symptomatic infected leaves four months after inoculating. Both isolates XYL2055/17 ($n$ = 316 inoculated plants) and XYL2177/18 ($n$ = 260) were collected from vineyards on Majorca. Scores were pooled among the 21 varieties inoculated;**(b)**conceptual graph of the population dynamics of *P. spumarius* on vineyards in Majorca and the effect on winter curing. Blue: density function of *P. spumarius*; red line: proportion of *P. spumarius* carrying Xf; blue line: proportion of plants recovering according the time they are infected;**(c)** bimodal density function of the number of symptomatic leaves. Blue dash line marks 5 symptomatic leaves; and **(d)**correlation between the upward and downward movement of $Xf_{PD}$ within the canes from the inoculation point. Each point depicts mean distance travelled in both directions by the bacteria.

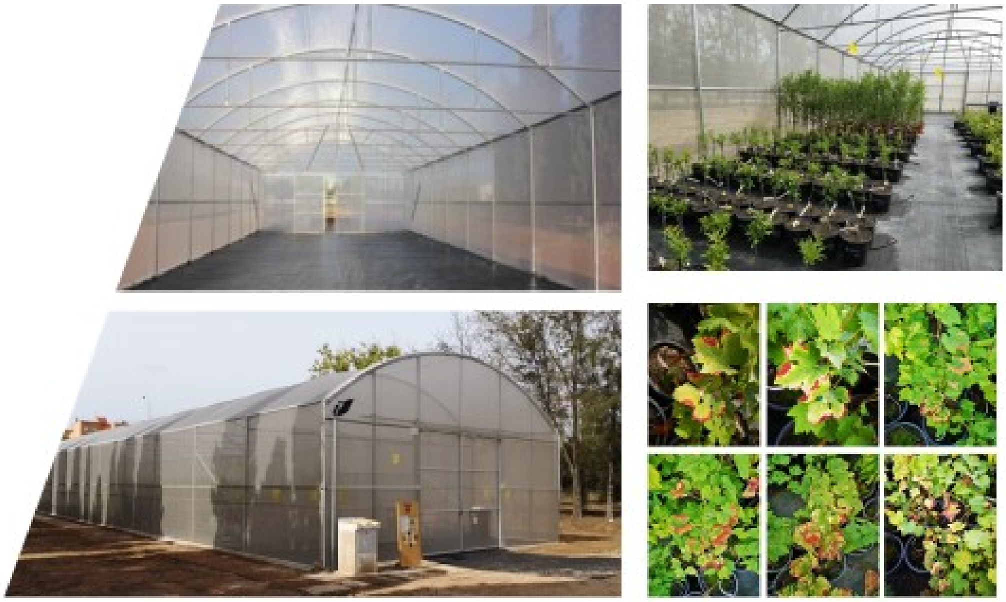

FIG. S2. **Experimental setup.** Greenhouse facilities and general view of its interior and the arrangement of the vine plants. The metallic structure is covered with an anti-thrips mesh.

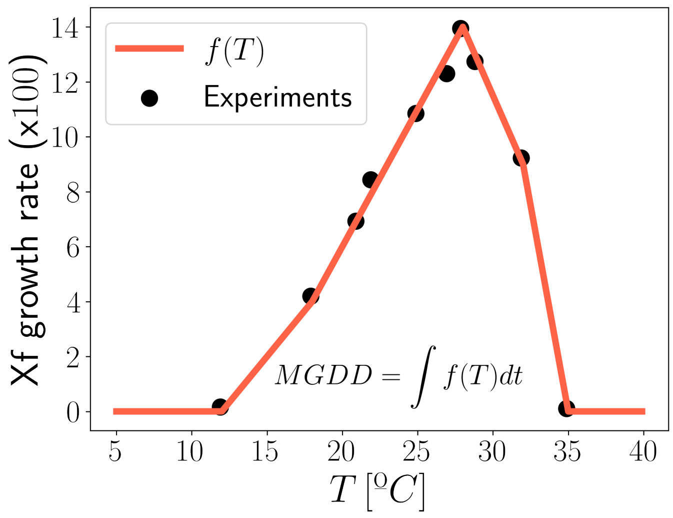

FIG. S3. **Relationship between MGDD and temperature.** Contribution to the $MGDD$ resulting from the fitting to the data in (1). The original Arrhenius plot, $\log k$ $vs.$ $1/T$ in kelvin was converted to a linear dependence in Celsius temperature $t$ (cf. Eq. (S3))

.

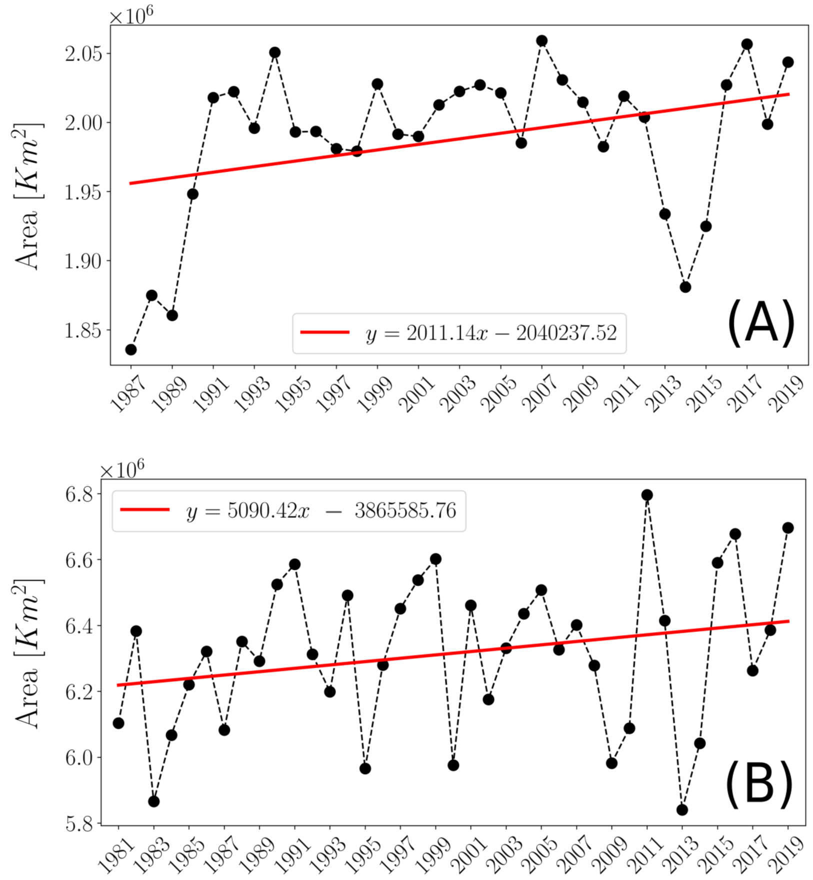

FIG. S4. **Trends in the risk-epidemic zones during the 1981-2019 period (A) and the areas encompassed below the $CDD < 314$ line (B) comprising land areas between $103^o$W and $70^o$W of the United States**

.

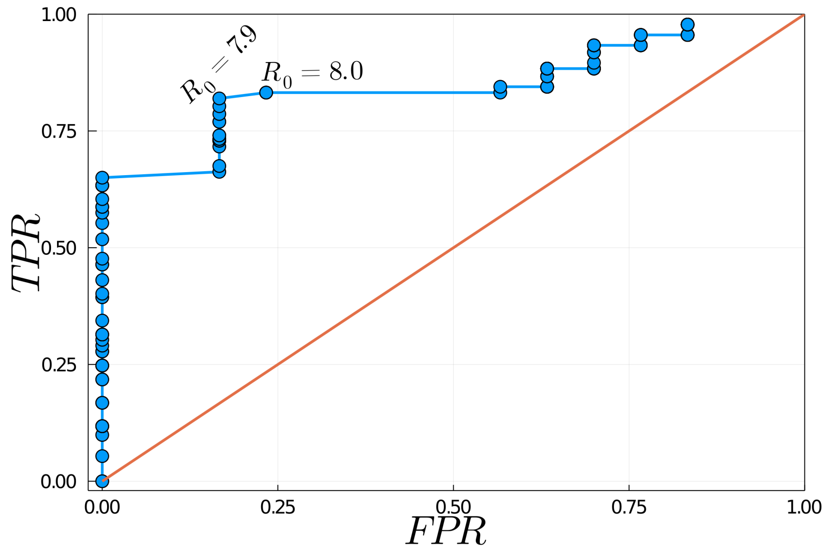

FIG. S5. **ROC curve illustrating the model validation procedure with spatiotemporal data from PD distribution in the US.** TPR is the *true positive rate* and FPR the *false positive rate*. The model accuracy reaches its optimum in $R_0 \approx 8$ by maximising the true positive rate and minimising the false positive rate. The spatiotemporal PD distribution in the US was obtained from data collected from publications between 2001 and 2015.

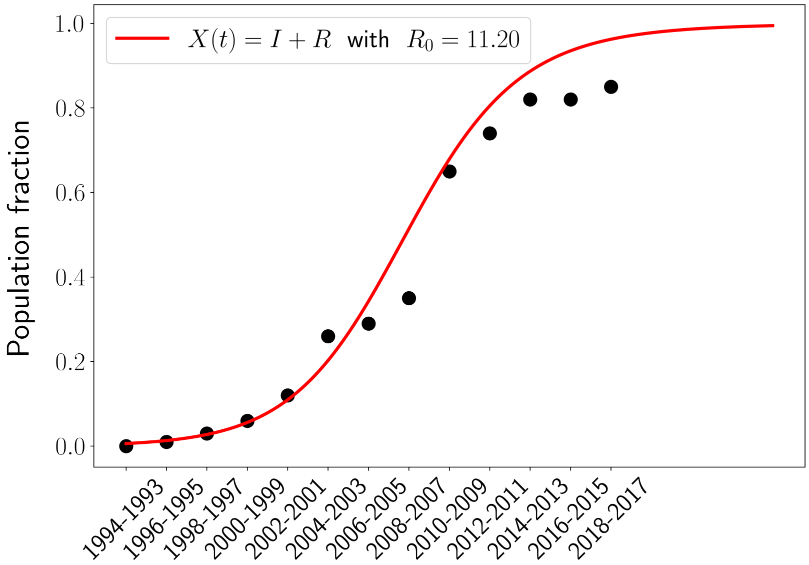

FIG. S6. **Fitting a $SIR$ model to the progress of the almond leaf scorch disease in the Balearic Islands from 1993 onward**.The best match was obtained with $R_0 = 11.2$. Points represent an estimate of the proportion of infected trees (incidence) from dendrochronological analysis and detection of Xf DNA in growth rings by qPCR. The incidence of ALSD in 2012 and 2017 was independently validated by field and Google Map Street View image observations.

.

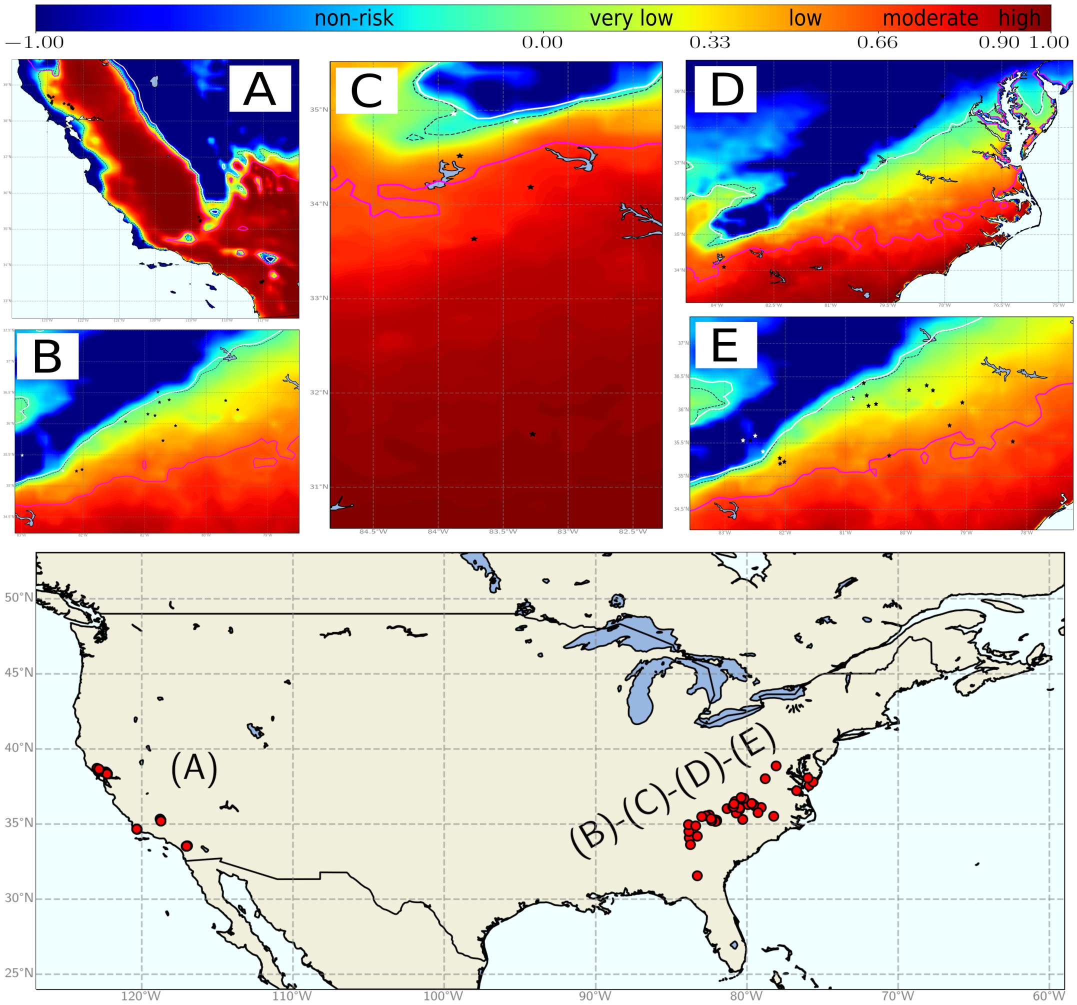

FIG. S7. **Model validation for an $R_0 = 8$ scenario with presence/absence data (black/white stars) of PD in the United States.** Panel (A) corresponds to data from California in 2015 while the other panels show data from 2002, 2005, 2006 and 2001 (respectively) in the east of the United States. The last panel clarifies the validation zones previously mentioned.

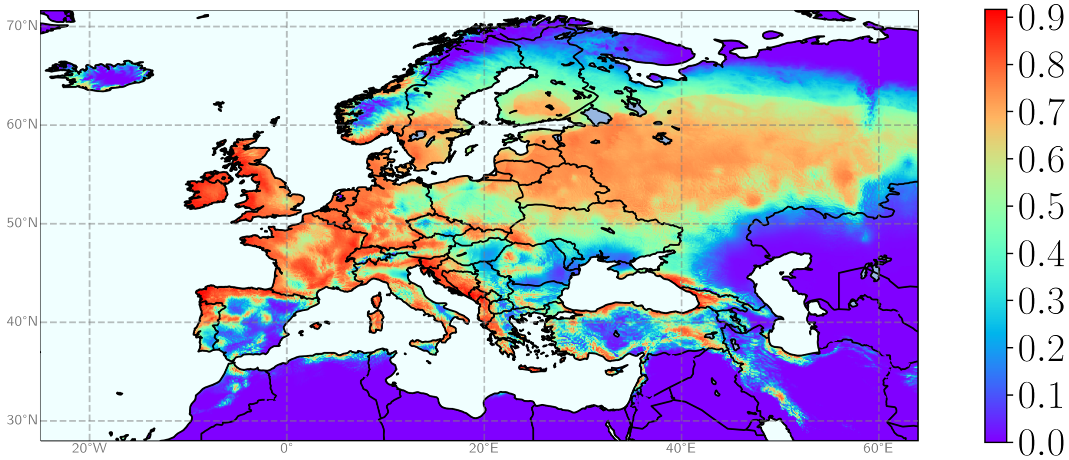

FIG. S8. **Average climatic suitability for *Philaenus spumarius* in Europe.** The map shows the climatic suitability of the vector estimated from a generalized additive model of insect distribution and the correlation of two bioclimatic descriptors, a climatic humidity index for the period of 8 coldest months of the year and the average maximum temperature in spring.

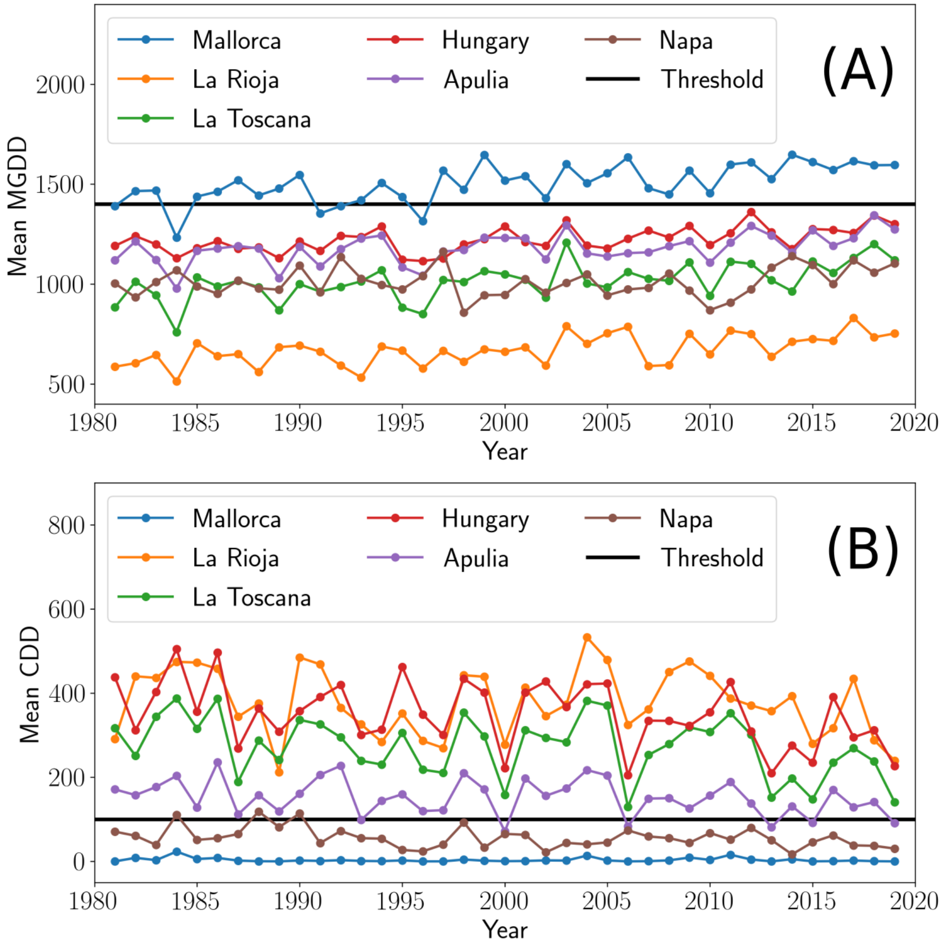

FIG. S9. **Trends in $MGDD$ (A) and $CDD$ (B) values and oscillations during 1981-2019 for seven wine regions with different climates from Europe and the US.** $MGDD$s show a slight upward trend and lesser oscillations than the $CDD$s.

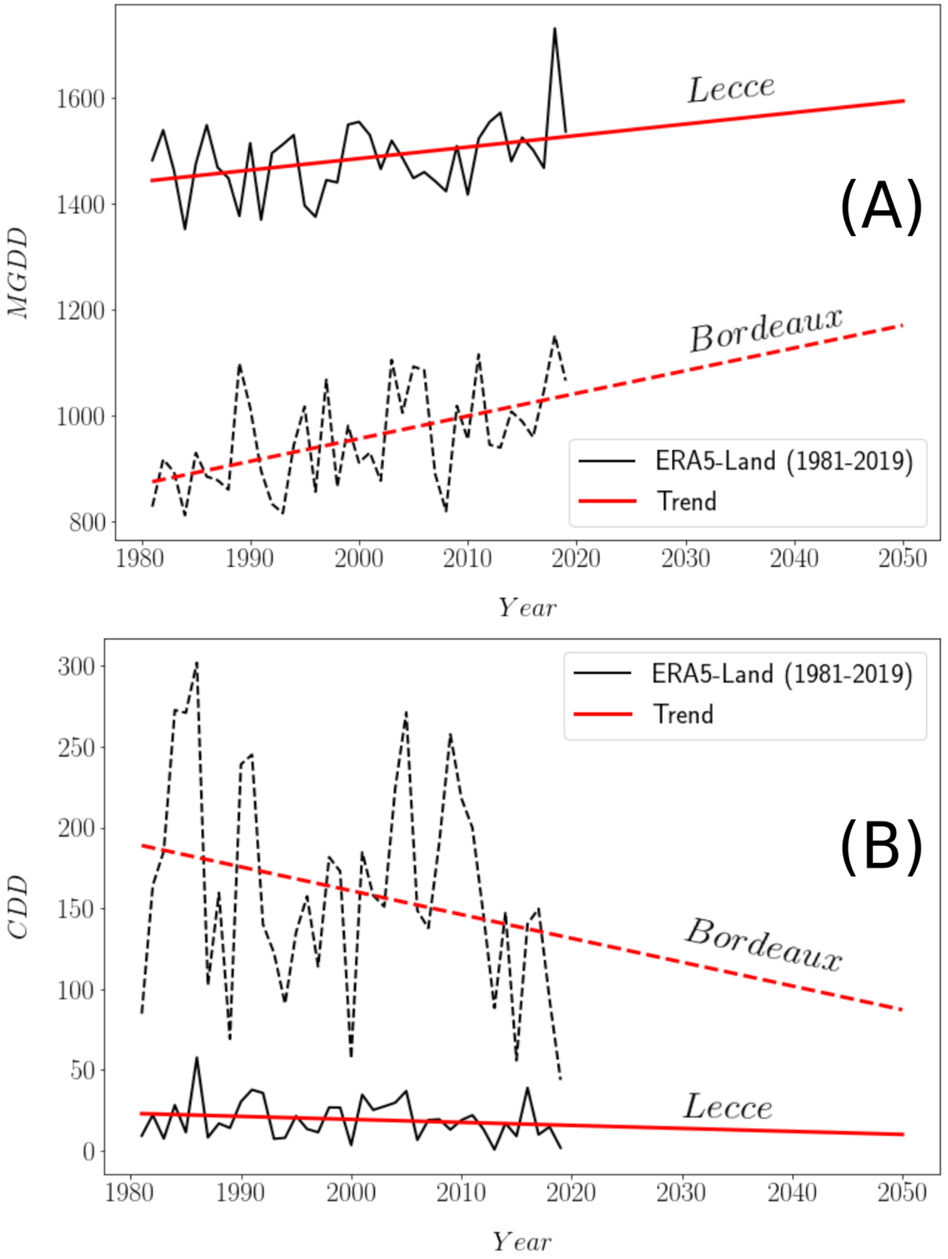

FIG. S10. **Determination of $MGDD$ and $CDD$ metric trends and future projections for two different European regions.** The $MGDD$ (A) and $CDD$ (B) trends show steeper slopes in the temperate climate of Bordeaux than in the Mediterranean climate of Lecce.

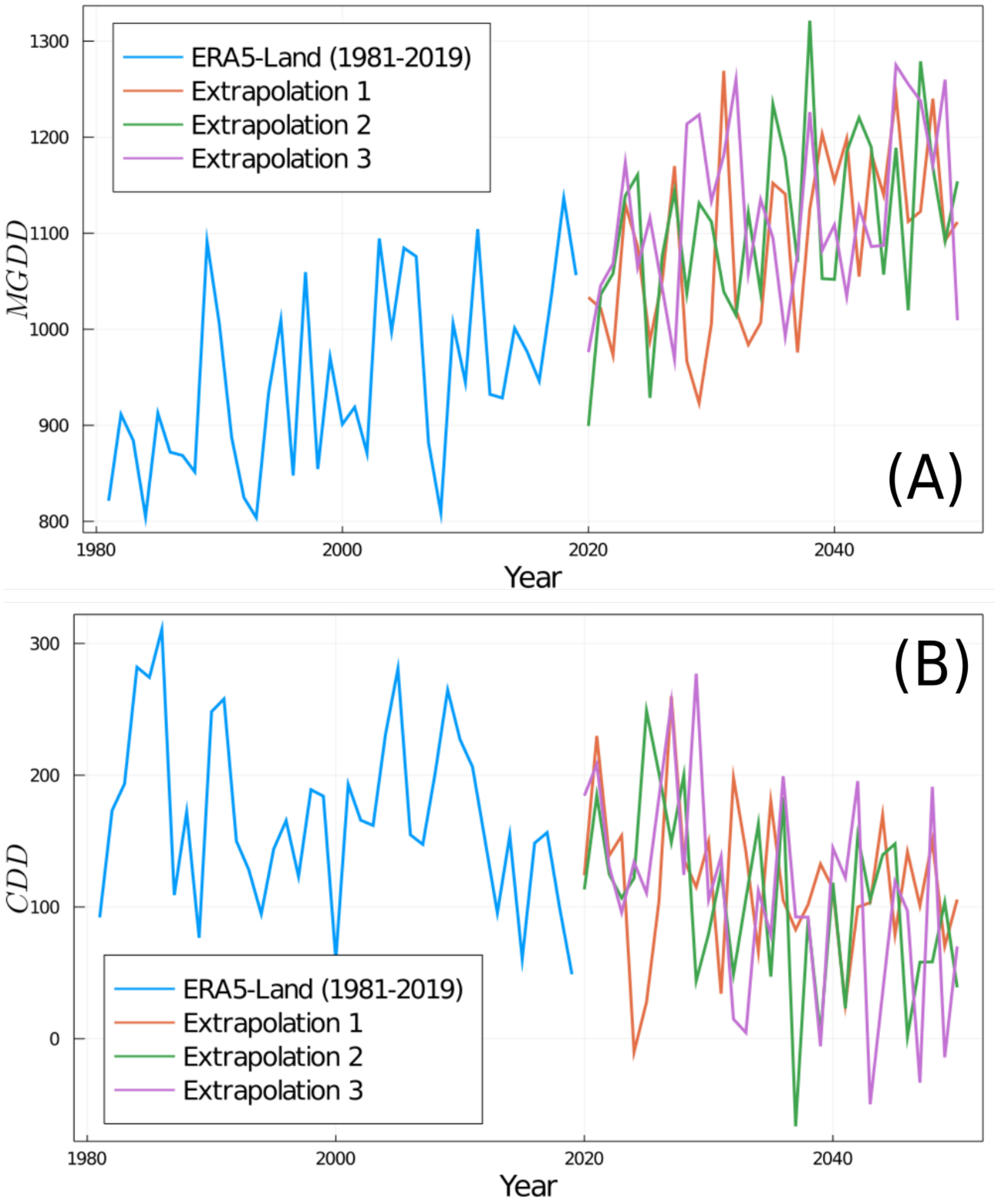

FIG. S11. **Interannual climatic variability extrapolations of** $MGDD$ **(A) and** $CDD$ **(B) for Bordeaux.** A linear model was fitted using Sklearn's LinearRegression module in Python and the interannual climatic variability was included as a Gaussian noise distribution by calculating the mean and fluctuations of the variance around $MGDD$ and $CDD$ trends.

### TABLES

TABLE S1: **Summary of the inoculation tests on grapevine varieties ranked from most to less susceptible in the disease index.** Thirty-six local, regional and international varieties were screened in combination with eight rootstocks. The number of symptomatic leaves were counted 16 weeks after inoculation and infections confirmed by qPCR. DI: disease index; AUDCP: area under the disease progress curve.

| Scion | Rootstock | Nº leaves | DI | AUDCP | % Positive | Year |
|---|---|---|---|---|---|---|
| Gorgollassa | R110 | 24.5 ± 8.8 | 5.00 | 31.29 | 100 | 2018 |
| Sauvignon Blanc | R110 | 16.5 ± 3.3 | 5.00 | 28.37 | 100 | 2018 |
| Tempranillo | SO4 | 15.7 ± 4.0 | 5.00 | 37.22 | 100 | 2019 |
| Garnacha tintorera | R110 | 19.0 ± 7.4 | 5.00 | 32.17 | 94.44 | 2019 |
| Tempranillo | 41B | 13.8 ± 2.7 | 4.89 | 13.43 | 0.00 | 2020 |
| Syrah | R140 | 13.6 ±3.0 | 4.86 | 29.46 | 98.21 | 2019 |
| Tempranillo Blanco | R110 | 16.0 ± 4.9 | 4.83 | 41.61 | 94.44 | 2019 |
| Chardonnay | R110 | 16.6 ± 5.6 | 4.83 | 26.39 | 94.44 | 2019 |
| Bobal | R110 | 15.2 ± 6.8 | 4.78 | 25.44 | 94.44 | 2019 |
| Prensal | 161/49 | 15.7 ± 5.0 | 4.75 | 21.37 | 100 | 2018 |
| Viura | SO4 | 16.7 ± 4.7 | 4.75 | 26.25 | 100 | 2018 |
| Garnacha tintorera | P1103 | 15.8 ± 6.8 | 4.72 | 35.94 | 94.44 | 2019 |
| Graciano | R140 | 14.3 ± 5.6 | 4.61 | 22.44 | 88.89 | 2019 |
| Airen | R110 | 15.2 ± 5.9 | 4.61 | 23.06 | 94.44 | 2019 |
| Mandó | R110 | 11 ± 3.6 | 4.56 | 21.22 | 100 | 2018 |
| Tempranillo | R110 RJ43 | 11.9 ± 3.0 | 4.56 | 31.17 | 100 | 2019 |
| Tempranillo | R110 RJ78 | 11.7 ± 3.8 | 4.50 | 34.33 | 94.44 | 2019 |
| Tempranillo | 41B | 12.4 ± 4.7 | 4.44 | 26.06 | 61.11 | 2019 |
| Garnacha | R110 | 11.1 ± 3.4 | 4.44 | 25.00 | 83.33 | 2019 |
| Viura | P1103 | 20.3 ± 8.4 | 4.37 | 25.875 | 87.5 | 2018 |
| Malvasia | R110 | 13.8 ± 6.5 | 4.33 | 26.71 | 71.73 | 2019 |
| Tempranillo | R110 RJ43 | 12.2 ± 5.9 | 4.33 | 24.23 | 88.89 | 2020 |
| Pedro Ximenez | R140 | 10.0 ± 4.33 | 4.33 | 16.87 | 0.00 | 2020 |
| Hondarrabi Beltza | 196-17 | 11.2 ± 4.6 | 4.22 | 19.90 | 87.5 | 2019 |
| Tempranillo | P1103 | 12.7 ± 6.3 | 4.17 | 30.62 | 62.5 | 2018 |
| Albariño | R110 | 9.5 ± 4.3 | 4.06 | 24.50 | 66.67 | 2019 |
| Garnacha tintorera | P1103 | 13.1 ± 8.0 | 4.0 | 16.85 | 88.89 | 2020 |
| Tempranillo | R110 | 11.7 ± 7.0 | 3.94 | 26.72 | 77.78 | 2019 |
| Merlot | R110 | 16.4 ± 13.1 | 3.75 | 24.87 | 75 | 2018 |
| Manto Negro | R110 | 11.9 ± 3.1 | 3.75 | 16.37 | | 2018 |
| Macabeo (Viura) | R110 | 13.0 ±9.6 | 3.72 | 19.54 | 70.83 | 2019 |
| Pinot Noir | R110 | 10.3 ± 7.1 | 3.67 | 13.42 | 44.44 | 2020 |
| Tempranillo Blanco | R110 | 11.2 ± 8.4 | 3.56 | 17.31 | 55.56 | 2020 |
| Syrah | R110 | 9.0 ± 5.6 | 3.50 | 12.37 | 100 | 2018 |
| Verdejo | R110 | 9.5 ± 6.6 | 3.50 | 13.89 | 72.22 | 2019 |
| Airen | R110 | 8.7 ± 5.3 | 3.44 | 15.98 | | 2020 |
| Monastrell | R110 | 12.3 ± 8.8 | 3.39 | 20.33 | 67.78 | 2019 |
| Mencia | R110 | 12.4 ± 9.8 | 3.39 | 18.28 | 61.11 | 2019 |
| Cabernet | R110 | 8.2 ± 6.0 | 3.33 | 11.56 | 72.22 | 2019 |
| Tempranillo | SO4 | 8.0 ± 4.9 | 3.33 | 20.44 | 44.44 | 2020 |
| Garnacha tintorera | R110 | 9.2 ± 7.6 | 3.33 | 15.68 | 55.56 | 2020 |
| Chardonnay | R110 | 7.0 ± 3.6 | 3.33 | 8.45 | 66.67 | 2020 |
| Viura | R140 | 10.5 ± 10.0 | 3.20 | 14.10 | 70 | 2018 |
| Pedro Ximénez | R110 | 7.5 ± 4.9 | 3.17 | 12.67 | 66.67 | 2019 |
| Garnacha | R110 | 9.2 ± 8.2 | 3.11 | 16.77 | 33.33 | 2020 |
| Pinot Noir | R110 | 8.4 ± 5.0 | 3.00 | 11.28 | 61.11 | 2019 |
| Cabernet Sauvignon | R110 | 6.6 ± 4.5 | 2.89 | 7.22 | 77.78 | 2020 |
| Syrah | R140 | 12.2 ± 10.0 | 2.89 | 15.44 | 55.56 | 2018 |
| Hondarrabi zuri | SO4 | 6.9 ± 6.3 | 2.78 | 12.78 | 61.11 | 2019 |
| Chardonnay | R110 | 8.4 ± 6.1 | 2.62 | 13.50 | 75 | 2018 |
| Graciano | R140 | 6.7 ± 5.7 | 2.56 | 8.12 | 55.56 | 2020 |
| Tempranillo | R140 | 8.1 ± 8.8 | 2.50 | 19.50 | 50 | 2018 |
| Tempranillo | R110 RJ78 | 5.3 ± 5.2 | 2.44 | 11.88 | 55.56 | 2020 |
| Prensal | R110 | 5.4 ± 5.5 | 2.37 | 9.25 | | 2018 |
| Tempranillo | SO4 | 9.7 ± 11.2 | 2.37 | 11.87 | 50 | 2018 |

| | | | | | | |
|---|---|---|---|---|---|---|
| Giró Ros | 161/49 | 6.1 ± 7.6 | 2.29 | 7.71 | | 2018 |
| Giró Negre | R110 | 5.2 ± 5.7 | 2.25 | 8.87 | 62.5 | 2018 |
| Viognier | R110 | 4.7 ± 5.0 | 2.25 | 6.50 | 75 | 2018 |
| Callet | R110 | 7.9 ± 9.8 | 2.25 | 8.87 | 37.5 | 2018 |
| Tempranillo | R110 | 3.9 ± 3.6 | 2.11 | 9.22 | 0.00 | 2020 |
| Hondarrabi beltza | 196-17 | 3.1 ± 1.8 | 1.89 | 5.89 | 11.11 | 2020 |
| Tempranillo | R110 | 4.0 ± 4.9 | 1.75 | 7.62 | 25 | 2018 |
| Argamussa | R110 | 3.2 ± 4.5 | 1.62 | 4.25 | | 2018 |
| Tempranillo | 41B | 5.6 ± 11.1 | 1.25 | 7.37 | 25 | 2018 |
| Albariño | R110 | 2.1 ± 2.1 | 1.22 | 5.23 | 33.33 | 2020 |
| Vinater Blanc | R110 | 2.0 ± 3.5 | 1.00 | 3.75 | 25 | 2018 |
| Hondarrabi zuri | SO4 | 1.6 ± 1.3 | 1 | 5.78 | 0.00 | 2020 |
| Cabernet | R110 | 2.9 ± 6.7 | 0.87 | 3.00 | 25 | 2018 |
| Syrah | 41B | 1.7 ± 3.9 | 0.87 | 3.75 | 12.5 | 2018 |
| Esperó de Gall | R110 | 4.0 ± 10.9 | 0.75 | 4.62 | 12.5 | 2018 |
| Sauvignon Blanc | SO4 | 0.5 ± 0.5 | 0.50 | 2.50 | 0 | 2018 |
| Giró Ros | R110 | 0.6 ± 1.9 | 0.37 | 1.50 | 0 | 2018 |
| Mancés | R110 | 0.4 ± 0.7 | 0.25 | 1.25 | | 2018 |

TABLE S2: **PD risk areas in Europe after running the model under a $R_0 = 5$ scenario and a homogeneous spatial vector distribution.** The epidemic-risk zones are classified according to the relative disease growth rates defined by the risk index, as very low, low, moderate and high growth rates. The total risk refers to the sum of the epidemic-risk zones

| **Country** | **No risk** ($km^2$) | **Transition** ($km^2$) | **Very low** ($km^2$) | **Low** ($km^2$) | **Moderate** ($km^2$) | **High** ($km^2$) | **Total risk** ($km^2$) | **Total surf.** ($km^2$) | **Risk** (%) |
|---|---|---|---|---|---|---|---|---|---|
| Russia | 4218776.2 | 15657.4 | 1698.2 | 0.0 | 0.0 | 0.0 | 1698.2 | 4236131.7 | 0.0 |
| Norway | 325425.8 | 0.0 | 0.0 | 0.0 | 0.0 | 0.0 | 0.0 | 325425.8 | 0.0 |
| France | 450678.5 | 36199.6 | 40474.0 | 8328.0 | 7015.8 | 1632.9 | 57450.7 | 544328.8 | 10.6 |
| Sweden | 441708.8 | 0.0 | 0.0 | 0.0 | 0.0 | 0.0 | 0.0 | 441708.8 | 0.0 |
| Belarus | 207565.9 | 0.0 | 0.0 | 0.0 | 0.0 | 0.0 | 0.0 | 207565.9 | 0.0 |
| Ukraine | 568981.5 | 0.0 | 0.0 | 0.0 | 0.0 | 0.0 | 0.0 | 568981.5 | 0.0 |
| Poland | 311522.7 | 0.0 | 0.0 | 0.0 | 0.0 | 0.0 | 0.0 | 311522.7 | 0.0 |
| Austria | 83265.0 | 0.0 | 0.0 | 0.0 | 0.0 | 0.0 | 0.0 | 83265.0 | 0.0 |
| Hungary | 92310.6 | 0.0 | 0.0 | 0.0 | 0.0 | 0.0 | 0.0 | 92310.6 | 0.0 |
| Moldova | 32139.5 | 0.0 | 0.0 | 0.0 | 0.0 | 0.0 | 0.0 | 32139.5 | 0.0 |
| Romania | 233239.3 | 1931.3 | 0.0 | 0.0 | 0.0 | 0.0 | 0.0 | 235170.6 | 0.0 |
| Lithuania | 64627.5 | 0.0 | 0.0 | 0.0 | 0.0 | 0.0 | 0.0 | 64627.5 | 0.0 |
| Latvia | 64206.0 | 0.0 | 0.0 | 0.0 | 0.0 | 0.0 | 0.0 | 64206.0 | 0.0 |
| Estonia | 45484.5 | 0.0 | 0.0 | 0.0 | 0.0 | 0.0 | 0.0 | 45484.5 | 0.0 |
| Germany | 354502.2 | 0.0 | 0.0 | 0.0 | 0.0 | 0.0 | 0.0 | 354502.2 | 0.0 |
| Bulgaria | 101668.8 | 8931.7 | 1277.8 | 0.0 | 0.0 | 0.0 | 1277.8 | 111878.3 | 1.1 |
| Greece | 37695.7 | 17152.2 | 17587.4 | 10467.4 | 14551.7 | 32116.9 | 74723.4 | 129571.2 | 57.7 |
| Albania | 18895.5 | 2417.9 | 2422.0 | 2045.2 | 2235.5 | 2142.2 | 8844.9 | 30158.3 | 29.3 |
| Croatia | 45699.9 | 2126.4 | 3268.6 | 1675.6 | 1059.9 | 270.0 | 6274.1 | 54100.3 | 11.6 |
| Switzerland | 46218.7 | 0.0 | 0.0 | 0.0 | 0.0 | 0.0 | 0.0 | 46218.7 | 0.0 |
| Luxembourg | 2705.7 | 0.0 | 0.0 | 0.0 | 0.0 | 0.0 | 0.0 | 2705.7 | 0.0 |
| Belgium | 30473.9 | 0.0 | 0.0 | 0.0 | 0.0 | 0.0 | 0.0 | 30473.9 | 0.0 |
| Netherlands | 36858.5 | 0.0 | 0.0 | 0.0 | 0.0 | 0.0 | 0.0 | 36858.5 | 0.0 |
| Portugal | 14211.9 | 14208.9 | 13810.1 | 5606.1 | 24768.7 | 15835.4 | 60020.2 | 88441.0 | 67.9 |
| Spain | 204150.9 | 41606.9 | 51545.3 | 53202.4 | 68116.9 | 85165.1 | 258029.7 | 503787.5 | 51.2 |
| Ireland | 68233.5 | 0.0 | 0.0 | 0.0 | 0.0 | 0.0 | 0.0 | 68233.5 | 0.0 |
| Italy | 112614.5 | 39537.2 | 46508.2 | 22088.7 | 30406.2 | 48759.8 | 147762.9 | 299914.6 | 49.3 |
| Denmark | 42273.9 | 0.0 | 0.0 | 0.0 | 0.0 | 0.0 | 0.0 | 42273.9 | 0.0 |
| United Kingdom | 242926.1 | 0.0 | 0.0 | 0.0 | 0.0 | 0.0 | 0.0 | 242926.1 | 0.0 |
| Iceland | 106696.1 | 0.0 | 0.0 | 0.0 | 0.0 | 0.0 | 0.0 | 106696.1 | 0.0 |
| Slovenia | 20237.3 | 258.0 | 86.4 | 0.0 | 0.0 | 0.0 | 86.4 | 20581.6 | 0.4 |
| Finland | 329375.9 | 0.0 | 0.0 | 0.0 | 0.0 | 0.0 | 0.0 | 329375.9 | 0.0 |
| Slovakia | 48140.8 | 0.0 | 0.0 | 0.0 | 0.0 | 0.0 | 0.0 | 48140.8 | 0.0 |
| Czechia | 80827.3 | 0.0 | 0.0 | 0.0 | 0.0 | 0.0 | 0.0 | 80827.3 | 0.0 |
| Bosnia and Herz. | 49554.0 | 449.7 | 0.0 | 0.0 | 0.0 | 0.0 | 0.0 | 50003.7 | 0.0 |
| Macedonia | 22838.2 | 2031.8 | 92.7 | 0.0 | 0.0 | 0.0 | 92.7 | 24962.7 | 0.4 |
| Serbia | 75992.9 | 0.0 | 0.0 | 0.0 | 0.0 | 0.0 | 0.0 | 75992.9 | 0.0 |
| Montenegro | 11374.8 | 363.7 | 455.1 | 729.9 | 365.7 | 0.0 | 1550.7 | 13289.1 | 11.7 |
| Kosovo | 11159.8 | 0.0 | 0.0 | 0.0 | 0.0 | 0.0 | 0.0 | 11159.8 | 0.0 |
| Cyprus | 404.2 | 0.0 | 0.0 | 0.0 | 101.1 | 4849.4 | 4950.5 | 5354.7 | 92.5 |
| Czech Republic | 78515.7 | 0.0 | 0.0 | 0.0 | 0.0 | 0.0 | 0.0 | 78515.7 | 0.0 |
| Malta | 0.0 | 0.0 | 0.0 | 0.0 | 0.0 | 199.6 | 199.6 | 199.6 | 100.0 |
| **TOTAL (%)** | 92.1 | 1.8 | 1.8 | 1.0 | 1.5 | 1.9 | 6.1 | | |

TABLE S3: **PD risk areas in the United States after running the model under a $R_0 = 8$ scenario and using a homogeneous spatial vector distribution.** The epidemic-risk zones are classified according to the relative disease growth rates defined by the risk index, as very low, low, moderate and high growth rates. The total risk refers to the sum of the epidemic-risk zones

| **State** | **No risk** (km$^2$) | **Transition** (km$^2$) | **Very low** (km$^2$) | **Low** (km$^2$) | **Moderate** (km$^2$) | **High** (km$^2$) | **Total risk** (km$^2$) | **Total surf.** (km$^2$) | **High Risk** (%) |
|---|---|---|---|---|---|---|---|---|---|
| Maine | 84556.5 | 0.0 | 0.0 | 0.0 | 0.0 | 0.0 | 0.0 | 84556.5 | 0.0 |
| Massachusetts | 20786.1 | 0.0 | 0.0 | 0.0 | 0.0 | 0.0 | 0.0 | 20786.1 | 0.0 |
| Michigan | 150179.3 | 0.0 | 0.0 | 0.0 | 0.0 | 0.0 | 0.0 | 150179.3 | 0.0 |
| Montana | 374475.7 | 0.0 | 0.0 | 0.0 | 0.0 | 0.0 | 0.0 | 374475.7 | 0.0 |
| Nevada | 239109.2 | 10675.3 | 7185.7 | 6918.9 | 10316.1 | 7978.1 | 32398.8 | 282183.3 | 2.8 |
| New Jersey | 15959.2 | 3802.3 | 286.4 | 0.0 | 0.0 | 0.0 | 286.4 | 20047.9 | 0.0 |
| New York | 127539.0 | 0.0 | 0.0 | 0.0 | 0.0 | 0.0 | 0.0 | 127539.0 | 0.0 |
| North Carolina | 15610.8 | 5709.3 | 16350.4 | 43864.7 | 40185.4 | 7502.2 | 107902.7 | 129222.9 | 5.8 |
| Ohio | 107590.9 | 0.0 | 0.0 | 0.0 | 0.0 | 0.0 | 0.0 | 107590.9 | 0.0 |
| Pennsylvania | 114772.4 | 0.0 | 0.0 | 0.0 | 0.0 | 0.0 | 0.0 | 114772.4 | 0.0 |
| Rhode Island | 2668.1 | 0.0 | 0.0 | 0.0 | 0.0 | 0.0 | 0.0 | 2668.1 | 0.0 |
| Tennessee | 3381.8 | 12419.7 | 65922.2 | 28656.9 | 0.0 | 0.0 | 94579.1 | 110380.6 | 0.0 |
| Texas | 3902.5 | 33646.7 | 35766.7 | 52625.6 | 196753.6 | 361569.4 | 646715.3 | 684264.5 | 52.8 |
| Utah | 211955.9 | 3627.2 | 1177.9 | 294.9 | 0.0 | 0.0 | 1472.9 | 217055.9 | 0.0 |
| Washington | 175209.0 | 171.5 | 0.0 | 0.0 | 0.0 | 0.0 | 0.0 | 175380.5 | 0.0 |
| Wisconsin | 144203.8 | 0.0 | 0.0 | 0.0 | 0.0 | 0.0 | 0.0 | 144203.8 | 0.0 |
| Puerto Rico | 1404.8 | 0.0 | 0.0 | 0.0 | 0.0 | 7724.3 | 7724.3 | 9129.1 | 84.6 |
| Maryland | 14568.2 | 5458.2 | 7044.3 | 291.0 | 0.0 | 0.0 | 7335.3 | 27361.8 | 0.0 |
| Alabama | 318.7 | 0.0 | 0.0 | 22524.9 | 41945.8 | 69646.2 | 134116.8 | 134435.5 | 51.8 |
| Alaska | 0.0 | 0.0 | 0.0 | 0.0 | 0.0 | 0.0 | 0.0 | 0.0 | 0.0 |
| Arizona | 89575.0 | 35856.7 | 16806.9 | 16919.7 | 37585.5 | 98303.0 | 169615.1 | 295046.9 | 33.3 |
| Arkansas | 0.0 | 8360.6 | 32714.2 | 46834.2 | 45982.8 | 0.0 | 125531.1 | 133891.7 | 0.0 |
| California | 130849.5 | 16184.8 | 22280.4 | 24793.2 | 61482.8 | 153642.1 | 262198.5 | 409232.7 | 37.5 |
| Colorado | 271791.7 | 0.0 | 0.0 | 0.0 | 0.0 | 0.0 | 0.0 | 271791.7 | 0.0 |
| Connecticut | 13262.1 | 0.0 | 0.0 | 0.0 | 0.0 | 0.0 | 0.0 | 13262.1 | 0.0 |
| Delaware | 950.0 | 2104.5 | 2308.4 | 0.0 | 0.0 | 0.0 | 2308.4 | 5362.9 | 0.0 |
| District of Columbia | 0.0 | 95.9 | 0.0 | 0.0 | 0.0 | 0.0 | 0.0 | 95.9 | 0.0 |
| Florida | 7163.0 | 0.0 | 0.0 | 0.0 | 0.0 | 142674.1 | 142674.1 | 149837.0 | 95.2 |
| Georgia | 525.5 | 202.1 | 3743.7 | 12387.4 | 35802.7 | 98902.3 | 150836.1 | 151563.7 | 65.3 |
| Hawaii | 0.0 | 0.0 | 0.0 | 0.0 | 0.0 | 0.0 | 0.0 | 0.0 | 0.0 |
| Idaho | 217685.6 | 0.0 | 0.0 | 0.0 | 0.0 | 0.0 | 0.0 | 217685.6 | 0.0 |
| Illinois | 136793.2 | 7722.9 | 0.0 | 0.0 | 0.0 | 0.0 | 0.0 | 144516.1 | 0.0 |
| Indiana | 91742.6 | 680.6 | 0.0 | 0.0 | 0.0 | 0.0 | 0.0 | 92423.1 | 0.0 |
| Iowa | 146768.4 | 0.0 | 0.0 | 0.0 | 0.0 | 0.0 | 0.0 | 146768.4 | 0.0 |
| Kansas | 202913.0 | 14336.5 | 0.0 | 0.0 | 0.0 | 0.0 | 0.0 | 217249.5 | 0.0 |
| Kentucky | 34786.1 | 58906.9 | 10265.3 | 0.0 | 0.0 | 0.0 | 10265.3 | 103958.3 | 0.0 |
| Louisiana | 6742.9 | 0.0 | 0.0 | 0.0 | 7966.1 | 108206.6 | 116172.7 | 122915.6 | 88.0 |
| Minnesota | 216457.3 | 0.0 | 0.0 | 0.0 | 0.0 | 0.0 | 0.0 | 216457.3 | 0.0 |
| Mississippi | 425.4 | 0.0 | 0.0 | 15007.2 | 54227.2 | 52624.1 | 121858.4 | 122283.8 | 43.0 |
| Missouri | 137772.8 | 36249.4 | 7829.0 | 0.0 | 0.0 | 0.0 | 7829.0 | 181851.2 | 0.0 |
| Nebraska | 194010.3 | 0.0 | 0.0 | 0.0 | 0.0 | 0.0 | 0.0 | 194010.3 | 0.0 |
| New Hampshire | 23784.7 | 0.0 | 0.0 | 0.0 | 0.0 | 0.0 | 0.0 | 23784.7 | 0.0 |
| New Mexico | 157196.9 | 31788.4 | 41306.7 | 41030.4 | 39489.5 | 0.0 | 121826.6 | 310812.0 | 0.0 |
| North Dakota | 180238.7 | 0.0 | 0.0 | 0.0 | 0.0 | 0.0 | 0.0 | 180238.7 | 0.0 |
| Oklahoma | 5031.2 | 45640.5 | 55077.9 | 58037.9 | 15414.7 | 0.0 | 128530.5 | 179202.2 | 0.0 |
| Oregon | 250298.0 | 601.0 | 0.0 | 0.0 | 0.0 | 0.0 | 0.0 | 250898.9 | 0.0 |
| South Carolina | 621.0 | 0.0 | 504.1 | 6161.4 | 38977.1 | 34104.7 | 79747.3 | 80368.3 | 42.4 |
| South Dakota | 200669.1 | 0.0 | 0.0 | 0.0 | 0.0 | 0.0 | 0.0 | 200669.1 | 0.0 |
| Vermont | 25033.8 | 0.0 | 0.0 | 0.0 | 0.0 | 0.0 | 0.0 | 25033.8 | 0.0 |
| Virginia | 44241.9 | 16964.7 | 28695.0 | 14762.2 | 787.6 | 0.0 | 44244.8 | 105451.4 | 0.0 |
| West Virginia | 62442.7 | 388.6 | 0.0 | 0.0 | 0.0 | 0.0 | 0.0 | 62831.4 | 0.0 |
| Wyoming | 252512.5 | 0.0 | 0.0 | 0.0 | 0.0 | 0.0 | 0.0 | 252512.5 | 0.0 |
| **TOTAl (%)** | 63.1 | 4.5 | 4.6 | 5.0 | 8.1 | 14.7 | 32.4 | | |

TABLE S4: **Potential distribution of PD in other world winegrowing regions.** In most areas of China and Australia *Vitis vinifera* is not cultivated and epidemic-risk zones with high growth rate correspond mainly to tropical areas in China, Australia, South Africa and Argentina

| **Country** | **No risk** ($km^2$) | **Transition** ($km^2$) | **Very low** ($km^2$) | **Low** ($km^2$) | **Moderate** ($km^2$) | **High** ($km^2$) | **Total risk** ($km^2$) | **Total surf.** ($km^2$) |
|---|---|---|---|---|---|---|---|---|
| *China* | 6,775,583.3 | 310,343.4 | 210,930.5 | 344,733.5 | 412,623.0 | 992,222.9 | 1,960,509.9 | 9,046,436.6 |
| *Australia* | 504,652.3 | 304,843.1 | 441,018.7 | 1,326,213.9 | 2,723,453.3 | 2,375,828.0 | 6,866,513.9 | 7,676,009.3 |
| *South Africa* | 216,593.4 | 183,726.3 | 119,839.8 | 152,000.0 | 279,105.5 | 264,223.2 | 815,168.6 | 1,215,488.2 |
| *Argentina* | 992,376.1 | 147,879.8 | 94,103.6 | 219,457.6 | 373,916.3 | 946,469.6 | 1,633,947.1 | 2,774,202.9 |
| *Chile* | 703,655.9 | 56,582.7 | 21,095.3 | 20,039.7 | 8,831.6 | 912.9 | 50,879.5 | 811,118.1 |

TABLE S5: **Predicted PD risk areas for the US in 2050 considering a $R_0 = 8$ scenario and a homogeneous spatial vector distribution.**The epidemic-risk zones are classified according to the relative disease growth rates defined by the risk index, as very low, low, moderate and high growth rates. The total risk refers to the sum of the epidemic-risk zones

| **State** | **No risk** (km$^2$) | **Transition** (km$^2$) | **Very low** (km$^2$) | **Low** (km$^2$) | **Moderate** (km$^2$) | **High** (km$^2$) | **Total risk** (km$^2$) | **Total surf.** (km$^2$) | **High Risk** (%) |
|---|---|---|---|---|---|---|---|---|---|
| Maine | 84556.5 | 0.0 | 0.0 | 0.0 | 0.0 | 0.0 | 0.0 | 84556.5 | 0.00 |
| Massachusetts | 20786.1 | 0.0 | 0.0 | 0.0 | 0.0 | 0.0 | 0.0 | 20786.1 | 0.00 |
| Michigan | 150179.3 | 0.0 | 0.0 | 0.0 | 0.0 | 0.0 | 0.0 | 150179.3 | 0.00 |
| Montana | 374475.7 | 0.0 | 0.0 | 0.0 | 0.0 | 0.0 | 0.0 | 374475.7 | 0.00 |
| Nevada | 230224.5 | 12349.0 | 8292.8 | 7785.0 | 10192.0 | 13339.9 | 39609.8 | 282183.3 | 4.73 |
| New Jersey | 10757.6 | 7004.1 | 2286.2 | 0.0 | 0.0 | 0.0 | 2286.2 | 20047.9 | 0.00 |
| New York | 126420.4 | 1118.7 | 0.0 | 0.0 | 0.0 | 0.0 | 0.0 | 127539.0 | 0.00 |
| North Carolina | 10300.2 | 4402.8 | 8608.6 | 28108.0 | 65840.0 | 11963.4 | 114520.0 | 129222.9 | 9.26 |
| Ohio | 107301.6 | 289.3 | 0.0 | 0.0 | 0.0 | 0.0 | 0.0 | 107590.9 | 0.00 |
| Pennsylvania | 114677.9 | 94.5 | 0.0 | 0.0 | 0.0 | 0.0 | 0.0 | 114772.4 | 0.00 |
| Rhode Island | 2668.1 | 0.0 | 0.0 | 0.0 | 0.0 | 0.0 | 0.0 | 2668.1 | 0.00 |
| Tennessee | 2287.5 | 1094.3 | 23757.5 | 79714.6 | 3526.7 | 0.0 | 106998.8 | 110380.6 | 0.00 |
| Texas | 2712.5 | 0.0 | 52099.2 | 46607.7 | 162037.0 | 420808.2 | 681552.0 | 684264.5 | 61.50 |
| Utah | 210192.6 | 2645.6 | 3333.5 | 785.8 | 98.4 | 0.0 | 4217.7 | 217055.9 | 0.00 |
| Washington | 165773.8 | 8239.3 | 1367.4 | 0.0 | 0.0 | 0.0 | 1367.4 | 175380.5 | 0.00 |
| Wisconsin | 144203.8 | 0.0 | 0.0 | 0.0 | 0.0 | 0.0 | 0.0 | 144203.8 | 0.00 |
| Puerto Rico | 1404.8 | 0.0 | 0.0 | 0.0 | 0.0 | 7724.3 | 7724.3 | 9129.1 | 84.61 |
| Maryland | 11612.4 | 4005.1 | 8649.1 | 3095.1 | 0.0 | 0.0 | 11744.2 | 27361.8 | 0.00 |
| Alabama | 318.7 | 0.0 | 0.0 | 3640.8 | 44508.9 | 85967.1 | 134116.8 | 134435.5 | 63.95 |
| Alaska | 0.0 | 0.0 | 0.0 | 0.0 | 0.0 | 0.0 | 0.0 | 0.0 | 0.00 |
| Arizona | 75250.5 | 27661.4 | 25822.3 | 14381.2 | 26927.3 | 125004.2 | 192135.0 | 295046.9 | 42.37 |
| Arkansas | 0.0 | 0.0 | 17518.8 | 41463.7 | 55669.6 | 19239.6 | 133891.7 | 133891.7 | 14.37 |
| California | 123859.6 | 9565.1 | 17917.7 | 21731.4 | 40312.5 | 195846.4 | 275808.0 | 409232.7 | 47.86 |
| Colorado | 254538.0 | 15192.6 | 2061.1 | 0.0 | 0.0 | 0.0 | 2061.1 | 271791.7 | 0.00 |
| Connecticut | 13262.1 | 0.0 | 0.0 | 0.0 | 0.0 | 0.0 | 0.0 | 13262.1 | 0.00 |
| Delaware | 381.4 | 854.1 | 3646.1 | 481.3 | 0.0 | 0.0 | 4127.4 | 5362.9 | 0.00 |
| District of Columbia | 0.0 | 95.9 | 0.0 | 0.0 | 0.0 | 0.0 | 0.0 | 95.9 | 0.00 |
| Florida | 7163.0 | 0.0 | 0.0 | 0.0 | 0.0 | 142674.1 | 142674.1 | 149837.0 | 95.22 |
| Georgia | 525.5 | 0.0 | 1011.1 | 4960.7 | 28065.0 | 117001.4 | 151038.2 | 151563.7 | 77.20 |
| Hawaii | 0.0 | 0.0 | 0.0 | 0.0 | 0.0 | 0.0 | 0.0 | 0.0 | 0.00 |
| Idaho | 213207.3 | 4478.4 | 0.0 | 0.0 | 0.0 | 0.0 | 0.0 | 217685.6 | 0.00 |
| Illinois | 118982.3 | 18687.9 | 6845.9 | 0.0 | 0.0 | 0.0 | 6845.9 | 144516.1 | 0.00 |
| Indiana | 84089.4 | 8333.7 | 0.0 | 0.0 | 0.0 | 0.0 | 0.0 | 92423.1 | 0.00 |
| Iowa | 146768.4 | 0.0 | 0.0 | 0.0 | 0.0 | 0.0 | 0.0 | 146768.4 | 0.00 |
| Kansas | 150938.1 | 45593.5 | 20717.9 | 0.0 | 0.0 | 0.0 | 20717.9 | 217249.5 | 0.00 |
| Kentucky | 15434.8 | 35258.9 | 52177.1 | 1087.4 | 0.0 | 0.0 | 53264.5 | 103958.3 | 0.00 |
| Louisiana | 6742.9 | 0.0 | 0.0 | 0.0 | 0.0 | 116172.7 | 116172.7 | 122915.6 | 94.51 |
| Minnesota | 216457.3 | 0.0 | 0.0 | 0.0 | 0.0 | 0.0 | 0.0 | 216457.3 | 0.00 |
| Mississippi | 425.4 | 0.0 | 0.0 | 505.3 | 43071.5 | 78281.7 | 121858.4 | 122283.8 | 64.02 |
| Missouri | 85361.3 | 56135.2 | 37967.8 | 2386.8 | 0.0 | 0.0 | 40354.7 | 181851.2 | 0.00 |
| Nebraska | 194010.3 | 0.0 | 0.0 | 0.0 | 0.0 | 0.0 | 0.0 | 194010.3 | 0.00 |
| New Hampshire | 23784.7 | 0.0 | 0.0 | 0.0 | 0.0 | 0.0 | 0.0 | 23784.7 | 0.00 |
| New Mexico | 136331.4 | 23902.7 | 42991.2 | 44481.8 | 51305.3 | 11799.5 | 150577.8 | 310812.0 | 3.80 |
| North Dakota | 180238.7 | 0.0 | 0.0 | 0.0 | 0.0 | 0.0 | 0.0 | 180238.7 | 0.00 |
| Oklahoma | 0.0 | 591.9 | 72476.3 | 69012.7 | 37121.2 | 0.0 | 178610.2 | 179202.2 | 0.00 |
| Oregon | 246416.5 | 2593.2 | 1889.3 | 0.0 | 0.0 | 0.0 | 1889.3 | 250898.9 | 0.00 |
| South Carolina | 621.0 | 0.0 | 0.0 | 1008.5 | 20253.1 | 58485.6 | 79747.3 | 80368.3 | 72.77 |
| South Dakota | 200669.1 | 0.0 | 0.0 | 0.0 | 0.0 | 0.0 | 0.0 | 200669.1 | 0.00 |
| Vermont | 25033.8 | 0.0 | 0.0 | 0.0 | 0.0 | 0.0 | 0.0 | 25033.8 | 0.00 |
| Virginia | 34490.9 | 11887.3 | 26642.0 | 30564.0 | 1867.2 | 0.0 | 59073.2 | 105451.4 | 0.00 |
| West Virginia | 56238.8 | 6592.6 | 0.0 | 0.0 | 0.0 | 0.0 | 0.0 | 62831.4 | 0.00 |
| Wyoming | 252512.5 | 0.0 | 0.0 | 0.0 | 0.0 | 0.0 | 0.0 | 252512.5 | 0.00 |
| **TOTAL (%)** | 59.6 | 4.0 | 5.6 | 5.2 | 7.6 | 18.1 | 36.5 | | |

TABLE S6: **Predicted PD risk areas in Europe in 2050 after running the model under a $R_0 = 5$ scenario and a homogeneous spatial vector distribution.** The epidemic-risk zones are classified according to the relative disease growth rates defined by the risk index, as very low, low, moderate and high growth rates. The total risk refers to the sum of the epidemic-risk zones

| **Country** | **No risk** ($km^2$) | **Transition** ($km^2$) | **Very low** ($km^2$) | **Low** ($km^2$) | **Moderate** ($km^2$) | **High** ($km^2$) | **Total risk** ($km^2$) | **Total surf.** ($km^2$) | **Risk** (%) |
|---|---|---|---|---|---|---|---|---|---|
| Russia | 4137884.7 | 60049.2 | 26738.7 | 10380.6 | 1078.6 | 0.0 | 38197.8 | 4236131.7 | 0.009 |
| Norway | 325425.8 | 0.0 | 0.0 | 0.0 | 0.0 | 0.0 | 0.0 | 325425.8 | 0.000 |
| France | 359832.7 | 62597.8 | 44497.8 | 53645.1 | 12403.9 | 11351.5 | 121898.3 | 544328.8 | 0.224 |
| Sweden | 441708.8 | 0.0 | 0.0 | 0.0 | 0.0 | 0.0 | 0.0 | 441708.8 | 0.000 |
| Belarus | 207565.9 | 0.0 | 0.0 | 0.0 | 0.0 | 0.0 | 0.0 | 207565.9 | 0.000 |
| Ukraine | 554548.4 | 13746.3 | 686.9 | 0.0 | 0.0 | 0.0 | 686.9 | 568981.5 | 0.001 |
| Poland | 311522.7 | 0.0 | 0.0 | 0.0 | 0.0 | 0.0 | 0.0 | 311522.7 | 0.000 |
| Austria | 81778.5 | 1486.5 | 0.0 | 0.0 | 0.0 | 0.0 | 0.0 | 83265.0 | 0.000 |
| Hungary | 39573.1 | 52737.6 | 0.0 | 0.0 | 0.0 | 0.0 | 0.0 | 92310.6 | 0.000 |
| Moldova | 32139.5 | 0.0 | 0.0 | 0.0 | 0.0 | 0.0 | 0.0 | 32139.5 | 0.000 |
| Romania | 206222.0 | 26493.7 | 2193.0 | 262.0 | 0.0 | 0.0 | 2455.0 | 235170.6 | 0.010 |
| Lithuania | 64627.5 | 0.0 | 0.0 | 0.0 | 0.0 | 0.0 | 0.0 | 64627.5 | 0.000 |
| Latvia | 64206.0 | 0.0 | 0.0 | 0.0 | 0.0 | 0.0 | 0.0 | 64206.0 | 0.000 |
| Estonia | 45484.5 | 0.0 | 0.0 | 0.0 | 0.0 | 0.0 | 0.0 | 45484.5 | 0.000 |
| Germany | 352649.4 | 1852.8 | 0.0 | 0.0 | 0.0 | 0.0 | 0.0 | 354502.2 | 0.000 |
| Bulgaria | 79172.5 | 22404.3 | 8386.3 | 1823.8 | 91.4 | 0.0 | 10301.5 | 111878.3 | 0.092 |
| Greece | 26086.7 | 9737.9 | 16221.2 | 17005.3 | 16275.7 | 44244.4 | 93746.6 | 129571.2 | 0.724 |
| Albania | 15635.2 | 2599.8 | 2987.2 | 2322.3 | 2885.1 | 3728.7 | 11923.4 | 30158.3 | 0.395 |
| Croatia | 22617.4 | 21932.1 | 2299.4 | 2657.7 | 2822.7 | 1770.9 | 9550.8 | 54100.3 | 0.177 |
| Switzerland | 46133.9 | 84.8 | 0.0 | 0.0 | 0.0 | 0.0 | 0.0 | 46218.7 | 0.000 |
| Luxembourg | 2705.7 | 0.0 | 0.0 | 0.0 | 0.0 | 0.0 | 0.0 | 2705.7 | 0.000 |
| Belgium | 30473.9 | 0.0 | 0.0 | 0.0 | 0.0 | 0.0 | 0.0 | 30473.9 | 0.000 |
| Netherlands | 36858.5 | 0.0 | 0.0 | 0.0 | 0.0 | 0.0 | 0.0 | 36858.5 | 0.000 |
| Portugal | 8015.9 | 5731.6 | 10805.1 | 12859.2 | 10320.1 | 40709.0 | 74693.5 | 88441.0 | 0.845 |
| Spain | 164633.8 | 40868.5 | 37919.6 | 54271.7 | 69026.9 | 137067.0 | 298285.2 | 503787.5 | 0.592 |
| Ireland | 68233.5 | 0.0 | 0.0 | 0.0 | 0.0 | 0.0 | 0.0 | 68233.5 | 0.000 |
| Italy | 83746.9 | 16519.8 | 31548.9 | 51473.6 | 38878.2 | 77747.1 | 199647.8 | 299914.6 | 0.666 |
| Denmark | 42273.9 | 0.0 | 0.0 | 0.0 | 0.0 | 0.0 | 0.0 | 42273.9 | 0.000 |
| United Kingdom | 242926.1 | 0.0 | 0.0 | 0.0 | 0.0 | 0.0 | 0.0 | 242926.1 | 0.000 |
| Iceland | 106696.1 | 0.0 | 0.0 | 0.0 | 0.0 | 0.0 | 0.0 | 106696.1 | 0.000 |
| Slovenia | 19211.5 | 596.6 | 429.2 | 258.0 | 86.4 | 0.0 | 773.5 | 20581.6 | 0.038 |
| Finland | 329375.9 | 0.0 | 0.0 | 0.0 | 0.0 | 0.0 | 0.0 | 329375.9 | 0.000 |
| Slovakia | 45501.4 | 2639.5 | 0.0 | 0.0 | 0.0 | 0.0 | 0.0 | 48140.8 | 0.000 |
| Czechia | 80827.3 | 0.0 | 0.0 | 0.0 | 0.0 | 0.0 | 0.0 | 80827.3 | 0.000 |
| Bosnia and Herz. | 44542.6 | 4741.3 | 719.9 | 0.0 | 0.0 | 0.0 | 719.9 | 50003.7 | 0.014 |
| Macedonia | 17314.7 | 4234.1 | 3228.6 | 185.4 | 0.0 | 0.0 | 3414.0 | 24962.7 | 0.137 |
| Serbia | 47601.5 | 28391.5 | 0.0 | 0.0 | 0.0 | 0.0 | 0.0 | 75992.9 | 0.000 |
| Montenegro | 10920.5 | 181.8 | 363.5 | 454.8 | 911.4 | 457.1 | 2186.8 | 13289.1 | 0.165 |
| Kosovo | 9705.9 | 1453.9 | 0.0 | 0.0 | 0.0 | 0.0 | 0.0 | 11159.8 | 0.000 |
| Cyprus | 404.2 | 0.0 | 0.0 | 0.0 | 0.0 | 4950.5 | 4950.5 | 5354.7 | 0.925 |
| Czech Republic | 78515.7 | 0.0 | 0.0 | 0.0 | 0.0 | 0.0 | 0.0 | 78515.7 | 0.000 |
| Malta | 0.0 | 0.0 | 0.0 | 0.0 | 0.0 | 199.6 | 199.6 | 199.6 | 1.000 |
| **TOTAL (%)** | 87.6 | 3.8 | 1.9 | 2.0 | 1.5 | 3.2 | 8.6 | | |

TABLE S7: **PD risk areas in Europe PD risk areas in Europe after running the model under a $R_0 = 5$ scenario and a spatial heterogeneous vector distribution (climatic suitability).** The epidemic-risk zones are classified according to the relative disease growth rates defined by the risk index, as very low, low, moderate and high growth rates. The total risk refers to the sum of the epidemic-risk zones

| **Country** | **No risk** ($km^2$) | **Transition** ($km^2$) | **Very low** ($km^2$) | **Low** ($km^2$) | **Moderate** ($km^2$) | **High** ($km^2$) | **Total risk** ($km^2$) | **Total surf.** ($km^2$) | **Risk (%)** |
|---|---|---|---|---|---|---|---|---|---|
| Russia | 4232704.4 | 3071.1 | 356.2 | 0.0 | 0.0 | 0.0 | 356.2 | 4236131.7 | 0.01 |
| Norway | 325425.8 | 0.0 | 0.0 | 0.0 | 0.0 | 0.0 | 0.0 | 325425.8 | 0.00 |
| France | 465951.2 | 57913.5 | 13241.5 | 6307.1 | 915.5 | 0.0 | 20464.1 | 544328.8 | 3.76 |
| Sweden | 441708.8 | 0.0 | 0.0 | 0.0 | 0.0 | 0.0 | 0.0 | 441708.8 | 0.00 |
| Belarus | 207565.9 | 0.0 | 0.0 | 0.0 | 0.0 | 0.0 | 0.0 | 207565.9 | 0.00 |
| Ukraine | 568981.5 | 0.0 | 0.0 | 0.0 | 0.0 | 0.0 | 0.0 | 568981.5 | 0.00 |
| Poland | 311522.7 | 0.0 | 0.0 | 0.0 | 0.0 | 0.0 | 0.0 | 311522.7 | 0.00 |
| Austria | 83265.0 | 0.0 | 0.0 | 0.0 | 0.0 | 0.0 | 0.0 | 83265.0 | 0.00 |
| Hungary | 92310.6 | 0.0 | 0.0 | 0.0 | 0.0 | 0.0 | 0.0 | 92310.6 | 0.00 |
| Moldova | 32139.5 | 0.0 | 0.0 | 0.0 | 0.0 | 0.0 | 0.0 | 32139.5 | 0.00 |
| Romania | 235170.6 | 0.0 | 0.0 | 0.0 | 0.0 | 0.0 | 0.0 | 235170.6 | 0.00 |
| Lithuania | 64627.5 | 0.0 | 0.0 | 0.0 | 0.0 | 0.0 | 0.0 | 64627.5 | 0.00 |
| Latvia | 64206.0 | 0.0 | 0.0 | 0.0 | 0.0 | 0.0 | 0.0 | 64206.0 | 0.00 |
| Estonia | 45484.5 | 0.0 | 0.0 | 0.0 | 0.0 | 0.0 | 0.0 | 45484.5 | 0.00 |
| Germany | 354502.2 | 0.0 | 0.0 | 0.0 | 0.0 | 0.0 | 0.0 | 354502.2 | 0.00 |
| Bulgaria | 110786.1 | 1092.2 | 0.0 | 0.0 | 0.0 | 0.0 | 0.0 | 111878.3 | 0.00 |
| Greece | 54287.5 | 27613.1 | 14844.7 | 17064.6 | 15761.4 | 0.0 | 47670.7 | 129571.2 | 36.79 |
| Albania | 19731.4 | 2885.8 | 2884.8 | 4561.5 | 94.8 | 0.0 | 7541.1 | 30158.3 | 25.01 |
| Croatia | 46319.6 | 2653.4 | 3094.4 | 1585.5 | 447.4 | 0.0 | 5127.3 | 54100.3 | 9.48 |
| Switzerland | 46218.7 | 0.0 | 0.0 | 0.0 | 0.0 | 0.0 | 0.0 | 46218.7 | 0.00 |
| Luxembourg | 2705.7 | 0.0 | 0.0 | 0.0 | 0.0 | 0.0 | 0.0 | 2705.7 | 0.00 |
| Belgium | 30473.9 | 0.0 | 0.0 | 0.0 | 0.0 | 0.0 | 0.0 | 30473.9 | 0.00 |
| Netherlands | 36858.5 | 0.0 | 0.0 | 0.0 | 0.0 | 0.0 | 0.0 | 36858.5 | 0.00 |
| Portugal | 19522.6 | 17715.8 | 45036.6 | 6166.1 | 0.0 | 0.0 | 51202.7 | 88441.0 | 57.89 |
| Spain | 262517.4 | 158650.5 | 66501.5 | 12133.5 | 3984.6 | 0.0 | 82619.6 | 503787.5 | 16.40 |
| Ireland | 68233.5 | 0.0 | 0.0 | 0.0 | 0.0 | 0.0 | 0.0 | 68233.5 | 0.00 |
| Italy | 133949.2 | 73040.4 | 41264.5 | 40374.0 | 11286.5 | 0.0 | 92924.9 | 299914.6 | 30.98 |
| Denmark | 42273.9 | 0.0 | 0.0 | 0.0 | 0.0 | 0.0 | 0.0 | 42273.9 | 0.00 |
| United Kingdom | 242926.1 | 0.0 | 0.0 | 0.0 | 0.0 | 0.0 | 0.0 | 242926.1 | 0.00 |
| Iceland | 106696.1 | 0.0 | 0.0 | 0.0 | 0.0 | 0.0 | 0.0 | 106696.1 | 0.00 |
| Slovenia | 20237.3 | 258.0 | 86.4 | 0.0 | 0.0 | 0.0 | 86.4 | 20581.6 | 0.42 |
| Finland | 329375.9 | 0.0 | 0.0 | 0.0 | 0.0 | 0.0 | 0.0 | 329375.9 | 0.00 |
| Slovakia | 48140.8 | 0.0 | 0.0 | 0.0 | 0.0 | 0.0 | 0.0 | 48140.8 | 0.00 |
| Czechia | 80827.3 | 0.0 | 0.0 | 0.0 | 0.0 | 0.0 | 0.0 | 80827.3 | 0.00 |
| Bosnia and Herz. | 49823.6 | 180.1 | 0.0 | 0.0 | 0.0 | 0.0 | 0.0 | 50003.7 | 0.00 |
| Macedonia | 24962.7 | 0.0 | 0.0 | 0.0 | 0.0 | 0.0 | 0.0 | 24962.7 | 0.00 |
| Serbia | 75992.9 | 0.0 | 0.0 | 0.0 | 0.0 | 0.0 | 0.0 | 75992.9 | 0.00 |
| Montenegro | 11465.8 | 363.7 | 728.9 | 730.8 | 0.0 | 0.0 | 1459.7 | 13289.1 | 10.98 |
| Kosovo | 11159.8 | 0.0 | 0.0 | 0.0 | 0.0 | 0.0 | 0.0 | 11159.8 | 0.00 |
| Cyprus | 404.2 | 0.0 | 1413.8 | 2223.9 | 1312.8 | 0.0 | 4950.5 | 5354.7 | 92.45 |
| Czech Republic | 78515.7 | 0.0 | 0.0 | 0.0 | 0.0 | 0.0 | 0.0 | 78515.7 | 0.00 |
| Malta | 0.0 | 0.0 | 0.0 | 0.0 | 199.6 | 0.0 | 199.6 | 199.6 | 100.00 |
| **TOTAL (%)** | 93.5 | 3.4 | 1.9 | 0.9 | 0.3 | 0.0 | 3.1 | | |

TABLE S8: **Surface of European vineyards in risk of PD given by the intersection of (Corine-Land-Cover) and the projected model in the ERA5-land data under a $R_0 = 5$ scenario with the layer of vector climatic suitability.** The epidemic-risk zones are classified according to the relative disease growth rates defined by the risk index, as very low (0.1-0.33), low (0.33-0.66), moderate (0.66-0.9) and high exponential growth rates ($> 90$). The total risk refers to the sum of the epidemic-risk zones.

| Country | No risk (Ha) | Transition (Ha) | Very low (Ha) | Low (Ha) | Moderate (Ha) | High (Ha) | Risk (%) |
|---|---|---|---|---|---|---|---|
| Albania | 1639.0 | 441.6 | 111.6 | 1370.8 | 0.0 | 0.00 | 41.61 |
| Austria | 66546.4 | 0.0 | 0.0 | 0.0 | 0.0 | 0.00 | 0.00 |
| Bulgaria | 112643.1 | 3203.0 | 0.0 | 0.0 | 0.0 | 0.00 | 0.00 |
| Switzerland | 14480.2 | 0.0 | 0.0 | 0.0 | 0.0 | 0.00 | 0.00 |
| Cyprus | 0.0 | 0.0 | 195.4 | 9535.4 | 4406.8 | 0.00 | 100 |
| Czech Republic | 16936.4 | 0.0 | 0.0 | 0.0 | 0.0 | 0.00 | 0.00 |
| Germany | 129648.9 | 0.0 | 0.0 | 0.0 | 0.0 | 0.00 | 0.00 |
| Greece | 13769.2 | 20611.6 | 13507.5 | 8536.7 | 24319.0 | 0.00 | 57.42 |
| Spain | 283550.1 | 696641.5 | 64083.4 | 5168.8 | 1211.1 | 0.00 | 6.71 |
| France | 371289.7 | 407458.9 | 282369.5 | 65302.9 | 3518.5 | 0.00 | 31.08 |
| Croatia | 16218.9 | 1519.1 | 3215.6 | 2578.3 | 1343.8 | 0.00 | 28.69 |
| Hungary | 100567.3 | 0.0 | 0.0 | 0.0 | 0.0 | 0.00 | 0.00 |
| Italy | 160369.5 | 180285.6 | 83666.8 | 116198.0 | 80098.6 | 0.00 | 45.11 |
| Luxembourg | 1633.9 | 0.0 | 0.0 | 0.0 | 0.0 | 0.00 | 0.00 |
| Montenegro | 0.0 | 1.9 | 2627.5 | 229.2 | 0.0 | 0.00 | 99.93 |
| Macedonia | 27731.9 | 0.0 | 0.0 | 0.0 | 0.0 | 0.00 | 0.00 |
| Malta | 25.7 | 0.0 | 0.0 | 0.0 | 27.4 | 0.00 | 51.54 |
| Portugal | 43689.1 | 74177.4 | 91311.8 | 2073.6 | 0.0 | 0.00 | 44.21 |
| Romania | 224664.7 | 0.0 | 0.0 | 0.0 | 0.0 | 0.00 | 0.00 |
| Serbia | 8610.0 | 0.0 | 0.0 | 0.0 | 0.0 | 0.00 | 0.00 |
| Slovenia | 25971.2 | 1173.9 | 840.6 | 0.0 | 0.0 | 0.00 | 3.00 |
| Slovakia | 20603.2 | 0.0 | 0.0 | 0.0 | 0.0 | 0.00 | 0.00 |
| **TOTAL (%)** | 42.1 | 35.6 | 13.9 | 5.4 | 3.0 | 0.00 | |

TABLE S9: **Predicted PD risk in 2050 in European vineyards (Corine-Land-Cover) considering a $R_0 = 5$ scenario and the vector climatic suitability.** The epidemic-risk zones are classified according to the relative disease growth rates defined by the risk index, as very low, low, moderate and high growth rates. The total risk refers to the sum of the epidemic-risk zones

| **Country** | **No risk (Ha)** | **Transition (Ha)** | **Very low (Ha)** | **Low (Ha)** | **Moderate (Ha)** | **High (Ha)** | **Risk (%)** |
|---|---|---|---|---|---|---|---|
| Albania | 766.7 | 921.5 | 475.7 | 1353.4 | 45.8 | 0.00 | 52.62 |
| Austria | 66546.4 | 0.0 | 0.0 | 0.0 | 0.0 | 0.00 | 0.00 |
| Bulgaria | 109593.3 | 5725.8 | 526.9 | 0.0 | 0.0 | 0.00 | 0.45 |
| Switzerland | 14480.2 | 0.0 | 0.0 | 0.0 | 0.0 | 0.00 | 0.00 |
| Cyprus | 0.0 | 0.0 | 848.6 | 12325.1 | 964.0 | 0.00 | 100.00 |
| Czech Republic | 16936.4 | 0.0 | 0.0 | 0.0 | 0.0 | 0.00 | 0.00 |
| Germany | 129648.9 | 0.0 | 0.0 | 0.0 | 0.0 | 0.00 | 0.00 |
| Greece | 13374.0 | 13093.0 | 24516.5 | 7907.9 | 21852.7 | 0.00 | 67.22 |
| Spain | 365465.7 | 617535.3 | 60443.8 | 5999.0 | 1211.1 | 0.00 | 6.44 |
| France | 245133.5 | 198142.5 | 510451.2 | 170502.6 | 5709.7 | 0.00 | 60.77 |
| Croatia | 15852.8 | 243.1 | 1880.0 | 6198.6 | 701.1 | 0.00 | 35.29 |
| Hungary | 100567.3 | 0.0 | 0.0 | 0.0 | 0.0 | 0.00 | 0.00 |
| Italy | 92799.8 | 131235.1 | 228546.2 | 155434.6 | 12602.7 | 0.00 | 63.90 |
| Luxembourg | 1633.9 | 0.0 | 0.0 | 0.0 | 0.0 | 0.00 | 0.00 |
| Montenegro | 0.0 | 0.0 | 1.9 | 2856.6 | 0.0 | 0.00 | 100.00 |
| Macedonia | 26532.8 | 1199.1 | 0.0 | 0.0 | 0.0 | 0.00 | 0.00 |
| Malta | 25.7 | 0.0 | 0.0 | 0.0 | 27.4 | 0.00 | 51.54 |
| Portugal | 19428.0 | 76697.5 | 101259.2 | 13867.1 | 0.0 | 0.00 | 54.50 |
| Romania | 224664.7 | 0.0 | 0.0 | 0.0 | 0.0 | 0.00 | 0.00 |
| Serbia | 8610.0 | 0.0 | 0.0 | 0.0 | 0.0 | 0.00 | 0.00 |
| Slovenia | 22469.8 | 1606.6 | 3068.7 | 840.6 | 0.0 | 0.00 | 13.97 |
| Slovakia | 20603.2 | 0.0 | 0.0 | 0.0 | 0.0 | 0.00 | 0.00 |
| **TOTAL (%)** | 38.4 | 26.9 | 23.9 | 9.7 | 1.1 | 0.0 | |

[17] M. Martcheva, *An Introduction to Mathematical Epidemiology* (Springer, New York, 2015).

[18] O. Diekmann, J. A. P. Heesterbeek, and M. G. Roberts, The construction of next-generation matrices for compartmental epidemic models, Journal of The Royal Society Interface **7**, 873 (2010).

[19] A. Giménez-Romero, R. Flaquer-Galmés, and M. A. Matias, Vector-borne diseases with non-stationary vector populations: the case of growing and decaying populations (2022).
